# Epithelial-Intrinsic Alterations and Maladaptation to Luminal Metabolites Underlie Persistent Crohn’s Disease Pathogenesis

**DOI:** 10.64898/2026.07.29.741535

**Authors:** Jianqiao Liu, Ran Zhou, Jason Koval, Peter Carbonetto, Candace M. Cham, Ashley M. Sidebottom, Matthew Stephens, Sebastian Pott, Eugene B. Chang, Anindita Basu

## Abstract

In patients with Crohn’s disease, the noninflamed intestinal epithelium exhibits inflammation-like transcriptional signatures that persist even during clinical remission. However, it is unclear whether this signature is due to sustained environmental triggers or to the epithelium’s intrinsic immunological adaptions. To disentangle environmental and cell-intrinsic effects on disease signatures, we used noninflamed intestinal tissue biopsies from patients with active and inactive Crohn’s disease, as well as control donors, to generate donor-matched organoids enriched for intestinal epithelial cells. We collected single-cell RNA-seq and single-cell ATAC-seq data from intestinal tissues, and from their matched organoids before and after stimulation with luminal metabolites of patients. This approach allowed us to distinguish between the effects of extrinsic factors and the intrinsic alterations that persist when cells are removed from their local environment. We found that organoid cells retained epithelial-autonomous disease phenotypes, even when derived from patients in remission, whereas signatures from tissue cells showed evidence of immune- and microbial-epithelial crosstalk. In contrast to healthy control organoids, organoids from Crohn’s disease patients exhibited selective activation of disease-associated transcriptional changes and environment-responsive inflammatory chromatin remodeling after exposure to patient-derived luminal metabolites. Overall, our study suggests that disease-associated epithelial states are shaped by both cell-intrinsic dysregulation and microenvironmental cues.

## Introduction

Crohn’s disease (CD) patients commonly experience chronic, relapsing transmural inflammation that affects specific segments of the GI tract, the terminal ileum and proximal colon.^1^ Although inflammation fluctuates with periods of flares and remissions, comprehensive single-cell atlases have revealed persistent molecular alterations in local tissue-specific resident cells, even under clinical remission.^2–6^ These observations suggest that cellular dysfunction may persist in spite of the absence of inflammation, therefore raising a critical question regarding the mechanisms that drive disease chronicity and relapse: are these molecular changes driven by persistent dysregulation intrinsic to the cells themselves, or by the sustained cues from the surrounding microenvironment?

A cell’s state is a complex phenotype that is regulated by both cell-intrinsic (e.g., genetic variants, epigenetic state) and cell-extrinsic influences (e.g., intercellular interactions, environmental modulators). The intestinal epithelium, a vital barrier and mediator between the host and microbiota, integrates both arms of regulation to maintain a resilient self-renewing tissue architecture and the fidelity of responses to a highly dynamic luminal environment.^7,8^ Both intrinsic and extrinsic factors are also relevant in understanding CD pathogenesis. For example, genome-wide association studies^9–13^ have uncovered hundreds of risk loci associated with genes regulating epithelial barrier integrity, autophagy, and immune sensing pathways, demonstrating the importance of cell-intrinsic susceptibility. Meanwhile, extrinsic factors such as microbial dysbiosis also modulate these epithelial cell states.^14–16^ Such interplay challenges our ability to isolate the contributions from various sources of regulation and thus identify direct mechanistic drivers of disease. Overcoming this challenge will require tractable model systems that can recapitulate patient-derived cell states in isolation and allow perturbations in a controlled context to dissect the pathways that underlie disease persistence and relapse.

Patient-derived organoids (PDOs) established from intestinal stem cells provide a powerful reductionist model for mechanistic studies of disease.^17^ These three-dimensional epithelial cultures retain the genetic background of the source tissue but are isolated from the influences of immune, stromal, and microbial cells. Benefiting from this isolation, disease-associated alterations that persist in culture can thus be attributed to cell-autonomous pathologies intrinsic to the epithelium.^18^ We applied this approach to isolate epithelial-intrinsic effects in CD by generating patient-matched single-cell profiles of gene expression and chromatin accessibility in PDOs and their corresponding ancestral tissue from CD and non-IBD donors. Sampling from both terminal ileum and ascending colon, we generated PDOs and subjected organoids and biopsied tissues to scRNA-seq and scATAC-seq. We demonstrated that CD PDOs presented cell-autonomous disease changes regardless of donor remission status, including proliferative defects, upregulation of IBD risk genes, and inflammation scores indicative of tissue disease burden; however, we also observed divergence in specific gene expression programs and chromatin regulatory elements linked to the immune and microbial crosstalk that are only present in tissues.

To investigate whether environmental influences might contribute to the persistence or reactivation of inflammatory epithelial states, we exposed PDOs to luminal metabolites derived from CD patients, representing a functional readout of the complex luminal milieu. Subsequent single-cell profiling of these stimulated cultures revealed a selective activation of disease-associated transcriptional alteration and environment-responsive inflammatory chromatin remodeling in CD PDOs. Collectively, these findings suggest that CD epithelial cells are epigenetically and transcriptionally primed for maladaptive responses to luminal stimuli, supporting that disease-associated epithelial states result from both cell-intrinsic dysregulation as well as microenvironmental cues.

## Results

### Single-cell ‘omics of PDOs and matched primary tissue

We collected 6 samples from CD patients in remission (CD noninf), 3 samples from adjacent non-inflamed sites of CD patients with active inflammation (CD adj), and 6 samples from non-IBD controls (Ctrl), totaling 6 samples from terminal ileum (TI) and 9 samples from ascending colon (AC). Each sample was processed in parallel for two purposes (Figure 1A): (i) single-cell isolation followed by scRNA-seq and scATAC-seq, generating primary tissue datasets as previously described;^3,19^ and (ii) establishment of matched patient-derived organoids (PDOs), cultured in progenitor-enriching expansion medium and subsequently profiled by scRNA-seq and scATAC-seq.

**Figure 1:**
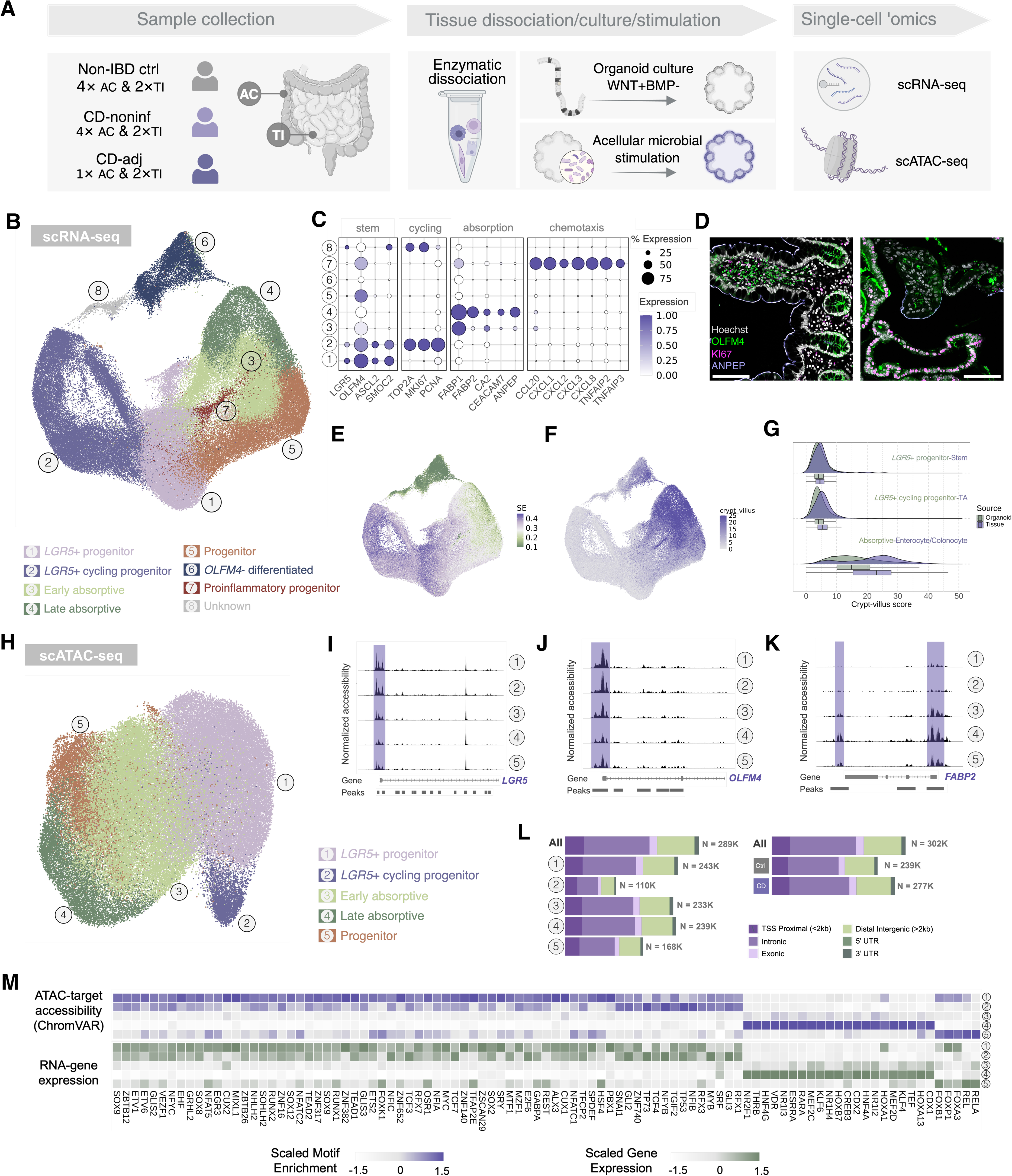
Multi-omics profiling of patient-derived organoids (PDOs). (A) Schematic illustration of the study design. (B) UMAP visualization of single-cell RNA sequencing (scRNA-seq) data from PDOs, colored by identified cell types. (C) A dot plot showing expression levels of selected marker genes across identified PDO cell types. (D) Immunofluorescence of cell type-specific markers in a tissue biopsy (left, patient HA56) and PDO (right, patient HA60) PDOs. Scale bar: 100 μm. (E) Signaling entropy scores of PDO cells. (F) Crypt-villus zonation scores of PDO cells. (G) Comparative analysis of crypt-villus differentiation scores between PDOs and matched tissue counterparts. (H) UMAP visualization of single-cell ATAC sequencing (scATAC-seq) data from PDOs, colored by identified cell types. (I–K) Coverage plots illustrating normalized chromatin accessibility at genomic peaks associated with marker genes for each cell type. (L) Distribution and quantification of accessible chromatin peaks identified within each cell type and disease condition, categorized by genomic region. (M) Top enriched transcription factor (TF) motifs identified from accessible chromatin regions (top) and expression levels of candidate regulatory TFs (bottom).

After quality control filtering, we recovered 41,528 epithelial transcriptomes from primary tissues (Figure S1A) and 102,849 transcriptomes from the corresponding PDOs (Figure 1B). Annotation of epithelial cells from primary tissues was guided by canonical marker genes (Figure S1B). We identified all major cell types previously reported for the absorptive and secretory epithelial lineages,^20,21^ including stem cells (*LGR5, OLFM4, ASCL2, SMOC2*), transit-amplifying (TA) cells (*MKI67, TOP2A*), ileal absorptive enterocytes (*FABP2, APOA4, ANPEP*), colonic absorptive colonocytes (*CA2*) and secretory cells, such as goblet cells (MUC2), BEST4 enterocytes (*BEST4*), Paneth cells (*DEFA5, DEFA6*), enteroendocrine cells (*CHGA*), and tuft cells (*POU2F3*).

Despite culture in media enriched with exogenous niche factors, PDOs exhibited spontaneous ‘differentiation’ into multiple epithelial lineages. As such, we applied a hybrid annotation strategy based on canonical markers and functional gene expression (Figure 1C). Among progenitor-like populations, we identified a classical *LGR5*⁺ progenitor state (*LGR5, OLFM4, ASCL2, SMOC2*) resembling stem cells from primary tissue, and a distinct proliferative *LGR5*⁺ *MKI67*⁺ population (*LGR5, MKI67, TOP2A*), which we termed *LGR5*⁺ cycling progenitors. Among more differentiated lineages, we annotated cells as early absorptive (*OLFM4, FABP1*), late absorptive (*FABP2, ANPEP*), and a proinflammatory progenitor state (*OLFM4, CCL20, CXCL*s, *TNFAIPs*). Notably, CCL20, also known as Macrophage inflammatory protein 3 alpha, is a chemokine historically linked to dome epithelial cells for the recruitment of *CCR6*⁺ immature dendritic cells^22,23^; its expression in this PDO lineage raises the intriguing possibility that this proinflammatory population may be relevant to gut-associated lymphoid tissue. A subset of cells lacking distinct lineage markers was categorized as Progenitor (*OLFM4*) or *OLFM4*-Differentiated. As expected, secretory lineages were absent in these expansion cultures, consistent with previous reports implicating p38 inhibitor (a component of the expansion media) in blocking secretory differentiation.^24,25^ We used immunofluorescence staining to confirm the expression of key markers including *OLFM4* (progenitors), *MKI67* (cycling progenitors), and *ANPEP* (late absorptive cells) in PDO cells of corresponding lineages as well as similar cell counterparts in tissues (Figure 1D).

To assess how these organoid cells order in terms of development trajectory and how they compared to the counterparts in primary tissue, we computed signaling entropy^26,27^ and crypt-villus score^28,29^. Signaling entropy is a proxy for cell potency, while crypt-villus score measures the progress of differentiating epithelial cells along the crypt-villus axis, with higher scores indicating later developmental stages. We observed a continuum of differentiation in both organoid and tissue (Figure 1E, 1F, Figure S1C, S1D), where progenitor populations exhibited higher signaling entropy and lower crypt-villus scores, while more differentiated states showed reduced potency and increased crypt-villus positioning, with a moderate correlation between two metrics (Spearman ρ=0.46, Figure S1E). Comparison across sources revealed similarly low crypt-villus scores in the stem and cycling cells from tissue and PDOs as a reflection of their undifferentiated states. However, the absorptive cells in organoid exhibited lower crypt-villus scores compared to those from tissue, suggesting an incomplete maturation *in vitro* (Figure 1G).

Regarding the scATAC-seq, we recovered 82,028 high-quality nuclei from PDOs (Figure 1H), and 23,901 epithelial nuclei from tissues (Figure S1F, S1G). In PDOs, we observed substantial patient-specific heterogeneity in chromatin accessibility (Figure S1H-S1J). To annotate the nuclei, we first performed data integration using reciprocal latent semantic indexing^30^ and then related the gene activity matrix inferred from scATAC-seq to the scRNA-seq count matrix. Chromatin accessibility information partly distinguished five major cell states in the PDOs (Figure 1H), forming tightly grouped clusters with minimal sample-to-sample variability after integration (Figure S1K). These included *LGR5*+ progenitor, *LGR5*+ cycling progenitor, early absorptive, late absorptive, and progenitor cells. These cell states were characterized by their differential accessibility at marker loci (Figure 1H-1J), including *LGR5*, *OLFM4*, and *FABP2*. In contrast, the epithelial nuclei from tissue samples showed broader diversity, reflecting a more complete *in vivo* differentiation landscape (Figure S1F).

To identify candidate *cis*-regulatory elements underlying PDO cell identity and disease-associated changes, we performed peak calling stratified by both cell type and disease condition (Figure 1L). This yielded a total of 288,565 unique peaks associated with cell state, with 109,744-242,859 peaks per state, and 301,867 unique peaks associated with disease condition, including 277,401 peaks specific to CD and 238,647 peaks specific to Ctrl. Genomic distribution of both peak sets was consistent and showed enrichment in enhancer-associated intronic and distal regions. Notably, *LGR5*⁺ cycling progenitors exhibited markedly fewer accessible peaks compared to other states, likely reflecting global chromatin compaction associated with mitosis.^31^ Overall, accessibility of cell-state-specific peaks corresponded closely with cell-type-specific gene expression. We combined all peaks into a union set comprising 319,179 regions for downstream analysis.

To predict upstream trans-regulators that putatively shape the chromatin accessibility landscape of various cell states and contributed to cell fate specification, we used chromVAR^32^ to compute the enrichment of transcription factor (TF) binding motifs among state-specific accessible peaks by comparing accessibility of motif-containing peaks to bias-matched background peaks. To increase confidence in predicting candidate regulators, we also calculated TF expression and its correlation with motif enrichment scores. We discovered 95 high-confidence candidate regulators (Figure 1M) that show strong correlations between TF motif enrichment and RNA-measured expression (Pearson ρ > 0.8). Many of the top-ranked TFs in organoids were consistent with established lineage-specific regulators in tissue, including *SOX9* in intestinal stem cells and *HNF4A*/*HNF4G* in absorptive enterocytes.

### CD PDOs harbor intrinsic proliferative defects

After data integration, we observed notable differences in cell-density distributions between CD and Ctrl organoids. Specifically, Ctrl PDOs were enriched for *LGR5*+ progenitors and cycling progenitors, whereas CD PDOs contained a higher proportion of more differentiated cells (Figure 2A, Figure S2A, S2B). To rigorously quantify disease-associated differences in cell-type composition, we employed a Dirichlet-multinomial regression model^33^. We grouped all cell types into three biologically meaningful categories, enabling a cross-source comparison between PDOs and matched primary tissues: (i) progenitors, which included *LGR5*+ progenitor and cycling progenitor in PDOs, and stem and TA in tissue, (ii) absorptive cells, which included early absorptive and late absorptive cells in PDOs, and enterocyte and colonocyte in tissue, and (iii) other cells, encompassing all remaining cell types. Our analysis revealed a significant decrease in the proportion of *LGR5*+ multipotent progenitors in PDOs derived from CD patients (p = 2.52 × 10^-2^, Figure 2B, 2C). No such reduction was observed in the corresponding primary tissues (p = 7.64 × 10^-1^, Figure S2C, S2D), suggesting this intrinsic defect can be masked by the native tissue environment. Interestingly, it has also been reported that PDOs derived from ulcerative colitis (UC) patients also demonstrated compromised regenerative potential,^34^ suggesting similarities in pathology of UC and CD.

**Figure 2:**
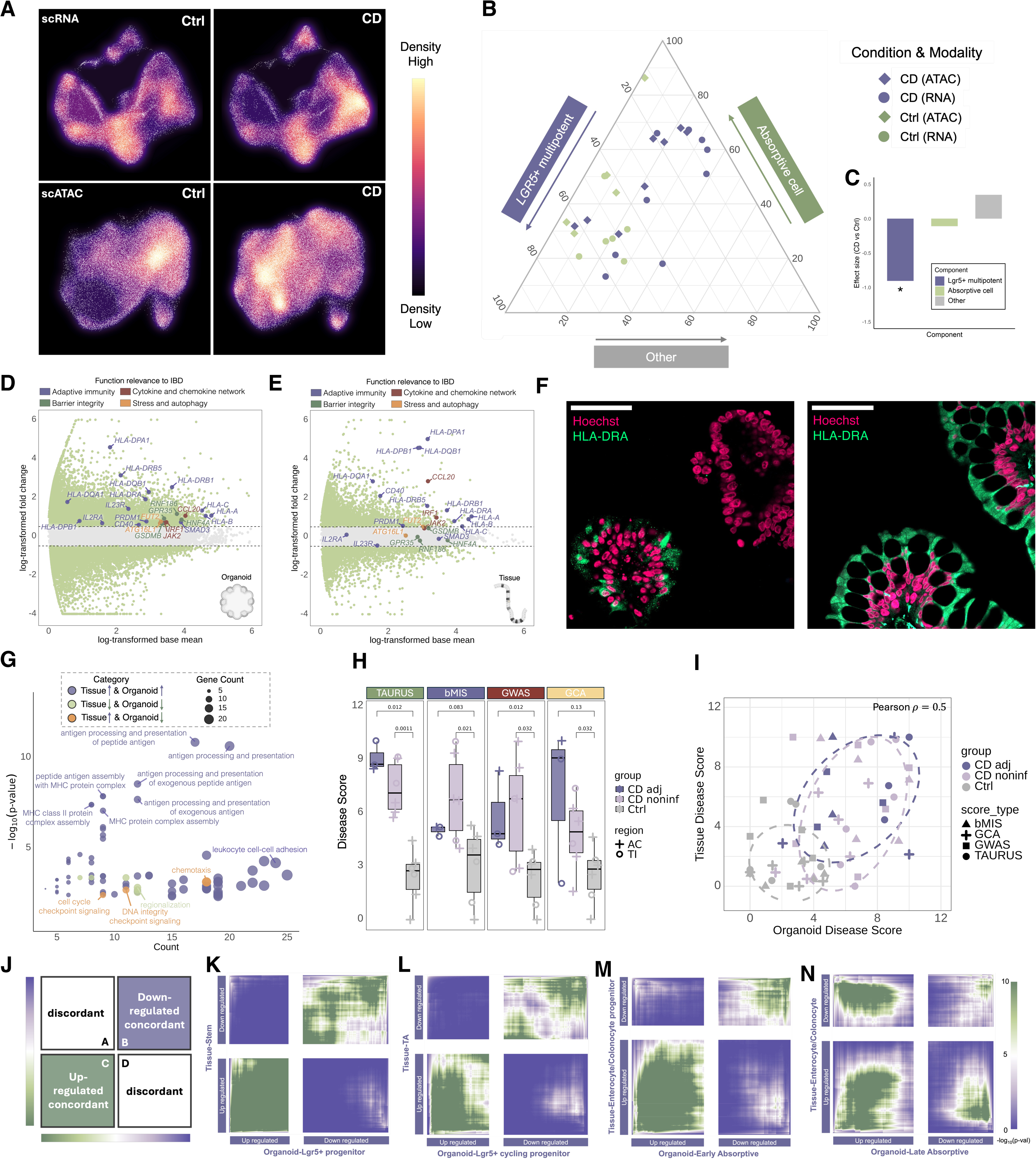
Cell-intrinsic alterations in CD PDOs and comparison to matched tissues. (A) UMAP visualization showing cell density distributions from scRNA-seq and scATAC-seq of PDOs. (B) A ternary plot illustrating relative cell-type compositions in PDO samples, with each datapoint representing an individual sample. (C) Dirichlet-multinomial regression analysis assessing cell composition changes specific to CD-derived PDOs. (D-E) MA plots displaying CD-specific differentially expressed genes (DEGs) identified in (D) PDOs and (E) matched tissues. (F) Immunofluorescence of HLA-DRA in a tissue biopsy (right, patient HA56) and matched PDO (left) demonstrating widespread expression in tissue versus sparse, heterogeneous expression in organoids. Scale bar: 50 μm. (G) A scatter plot comparing gene ontology enrichment results of DEGs between PDOs and matched tissues. (H) Inflammatory disease scoring of PDOs calculated using curated inflammatory gene sets. (I) Pearson correlation analysis of inflammatory disease scores between PDOs and matched tissues. (J) Schematic illustration of Rank-Rank Hypergeometric Overlap (RRHO2) plots; genes from each comparison are ordered from most upregulated to most downregulated, with the highest-ranked genes located in the lower-left quadrant. (K–N) RRHO2 plots comparing transcriptional signatures of corresponding cell types from matched PDOs and tissue samples. Heatmap color indicates the significance (–log₁₀ p-value) of gene-list overlap determined by hypergeometric tests.

### PDOs show both conserved and divergent expression profiles for core IBD risk genes

After analyzing cell composition differences in tissues and matched PDOs, we sought to determine whether and what disease-specific transcriptional signatures were preserved in PDOs. To that end, we performed differentially expressed gene (DEG) analysis, comparing expression levels between disease and control conditions in tissues and PDOs. We found that many core IBD risk genes were consistently upregulated in both tissue and PDOs from CD patients (Table 1, Figure 2D, 2E), comprising three distinct categories of functional relevance^13^ to IBD. MHC-mediated adaptive immunity (*HLA-A*, *HLA-B*, *HLA-C*, *HLA-DRB1*, *HLA-DRA*, *HLA-DRB5*, *HLA-DPA1*, *HLA-DPB1*, *HLA-DQB1*, *HLA-DQA1*), cytokine networks (*CCL20, IRF1*), and stress response (*FUT2*). This transcriptional signature was observed in both CD noninf and CD adj samples, albeit with some differences in magnitude and composition (Figure S2E-S2H). These genes represented persistent molecular alterations independent of environmental triggers, reflecting cell-intrinsic programs that may contribute to chronic disease susceptibility or relapse.

**Table 1:** Differential expression of core IBD risk genes in organoid and tissue and their functional relevance to mucosal immunity.

| Gene name | Functional relevance | IBD risk pathways | Differential expression source |
| --- | --- | --- | --- |
| *HLA* locus | Adaptive immunity | Antigen presentation | Organoid, Tissue |
| *PRDM1* | Adaptive immunity | T<sub>reg</sub> | Organoid, Tissue |
| *CD40* | Adaptive immunity | IgA | Organoid, Tissue |
| *CCL20* | Cytokine and chemokine network | Cytokine | Organoid, Tissue |
| *IRF1* | Cytokine and chemokine network | Cytokine | Organoid, Tissue |
| *FUT2* | Stress and autophagy | Granule biogenesis | Organoid, Tissue |
| *HNF4A* | Epithelial barrier integrity | Unfolded protein response | Organoid |
| *GPR35* | Epithelial barrier integrity | Barrier function | Organoid |
| *GSDMB* | Epithelial barrier integrity | IgA | Organoid |
| *RNF186* | Epithelial barrier integrity | Barrier function | Organoid |
| *SMAD3* | Adaptive immunity | T<sub>reg</sub> | Organoid |
| *IL2RA* | Adaptive immunity | T<sub>reg</sub> | Organoid |
| *IL23R* | Adaptive immunity | T<sub>h</sub>17 | Organoid |
| *ATG16L1* | Stress and autophagy | Autophagy | Organoid |
| *JAK2* | Cytokine and chemokine network | Cytokine | Organoid |
| *NOS2* | Stress and autophagy | Oxidative stress | Tissue |
| *NOD2* | Microbe sensing | Microbe sensing | Tissue |

While the MHC-II genes were transcriptionally upregulated in CD PDOs, their expression patterns differed in tissues and PDOs. In IBD tissues, intestinal epithelial cells constitutively express high levels of MHC-II genes,^33,35,36^ as confirmed by the widespread immunofluorescent staining of HLA-DRα in tissue epithelium (Figure 2F, Figure S2I). In contrast, the expression of HLA-DRα in PDOs was found to be heterogeneously sparse, in both scRNA-seq and immunofluorescence data (Figure 2F, Figure S2J).

We also identified some IBD risk genes that were DEGs exclusively in PDOs, and not detected as DEGs in matched tissues (Table 1). Their functional connections to IBD include epithelial barrier integrity (*HNF4A*, *GPR35*, *GSDMB*, *RNF186*), adaptive immunity (*IL2RA*, *IL23R*, *SMAD3*), and autophagy (*ATG16L1*). The selective upregulation of these genes in PDOs implies intrinsic liabilities in barrier maintenance, stress response, and adaptive immune crosstalk that are normally masked by complex interactions present within intact tissue microenvironments.

Contrasting DEGs between PDOs and tissues revealed 535 genes concordantly upregulated in both systems, while 333 genes specifically upregulated in tissue, and 459 genes uniquely upregulated in PDOs (Figure S2I). Gene Ontology enrichment analysis (Figure 2G) showed that concordantly upregulated genes were significantly enriched in processes related to the MHC protein complex assembly, antigen presentation, and leukocyte recruitment. Tissue-specific DEGs were primarily associated with cell cycle checkpoint signaling, DNA integrity checkpoint responses, and chemotaxis, whereas genes specifically upregulated in PDOs exhibited limited enrichment, notably in regionalization processes.

### Cell-autonomous CD signatures quantitatively correlate with epithelium disease burden

Previous studies have found that macroscopically noninflamed tissue can still harbor transcriptional signatures indicative of inflammation,^2,3,33^ underscoring the need for transcriptome-based inflammation scoring methods. Therefore, we asked whether PDOs could quantitatively recapitulate the inflammatory states of their corresponding ancestral tissues at the transcriptional level. We leveraged four different scoring feature lists from four different sources: the TAURUS study by Thomas *et al*.^5^ (TAURUS), biopsy-based biomarker of inflammation study^37^ (bMIS), genome-wide association study^38^ (GWAS), and our previous atlas study^3^ of Crohn’s disease (GCA). However, applying existing inflammation gene signatures directly to PDOs introduced a conceptual and practical challenge: these signatures were primarily derived from whole mucosal biopsy samples encompassing not only epithelial but also immune and stromal compartments, while PDOs only consisted of epithelial population in isolation. In addition, our PDOs were cultured in the undifferentiating expansion medium and thus represent only a subset of the functional spectrum within in the primary epithelium. Therefore, we refined the inflammation scoring features by intersecting established tissue-derived inflammation gene lists with the set of differentially expressed genes specifically identified in PDOs, generating an epithelium-centric gene panel suitable for evaluating inflammatory states intrinsic to epithelium (Table S2). Rather than defining disease status *de novo*, this panel was used to evaluate the extent to which PDOs recapitulate tissue-derived inflammatory signatures. Using this approach, we observed that inflammation scores derived from this panel stratified CD and non-IBD samples in both PDOs and matched tissues (Figure 2H, S2J). Importantly, Pearson correlation analysis revealed a moderate correlation (ρ=0.5) between the inflammation scores in PDOs and those in matched ancestral tissues. Overall, these results highlight that PDOs partially reflected the inflammatory status of their corresponding tissues at the transcriptional level and that the cell-intrinsic features they exhibited represented a quantitative measure of the overall disease burden (Figure 2I).

### PDO-tissue transcriptional concordance varies by differentiation stage

In the above analyses, we identified extensive overlap between the molecular signatures of CD in organoids and those defined in matched tissues. We next explored whether these similarities extend equally to all cell types across developmental stages, by applying a pairwise comparison of disease signatures defined separately for each pair of cell type counterparts in PDOs and tissues using rank-rank hypergeometric overlap^39,40^ (RRHO, Figure 2J). We found strong concordance between systems in these multipotent progenitor cells (Figure 2K, 2L)-i.e., between *LGR5*+ progenitor (PDO) and stem cells (tissue), as well as the *LGR5*+ cycling progenitor of PDOs and TA cells of tissue. These findings suggest that disease-specific transcriptional changes during early differentiation trajectories may be driven in large part by cell-intrinsic factors. However, notable discordance emerged in the absorptive cells (Figure 2M, 2N) with reduced overlap in disease-related signatures between early and late absorptive cells in PDOs and their respective enterocyte and colonocyte counterparts in tissue. This divergence was consistently observed in both CD non-inflamed and adjacent samples (Figure S2M-S2T), and suggests that the extracellular environment may contribute more than cell-autonomous regulation to disease-linked transcriptional changes for epithelial cells in later developmental stages.

### CD-linked gene expression programs in PDOs reflect the absence of immune and microbial crosstalk in PDOs

Partitioning cells into discrete cell states may not capture the full spectrum of cell identity and activity. Therefore, we leveraged non-negative matrix factorization^41–44^ (NMF), which extracts latent gene expression programs^45–47^ (GEPs), and then represents the cells as combinations of the GEPs. NMF identifies a set of paired vectors, one capturing the degree to which the GEP is active in each cell (“cell membership”), and the other defining the log-fold change in expression for each gene. This approach allowed us to decompose biological heterogeneity into modular, interpretable patterns of gene expression.

Applying NMF to the datasets from tissues and PDOs revealed GEPs distinct to tissues and PDOs, as well as a subset of similar GEPs that were similar in tissues and PDOs. This subset of similar GEPs included GEPs associated with CD pathology, cell cycling, and absorption (Figure 3A, 3B, S3A, S3B). We defined “pathology-relevant GEPs” based on strong membership in samples derived from CD patients and significant overlap of driving genes with known GWAS-identified CD risk loci (Figure S3C, S3D). Comparative analyses using a combined similarity measure (Jaccard index × cosine similarity) revealed several GEP modules shared between PDOs and tissues (Figure 3C). Cell-cycle phases predicted independently from canonical marker genes matched closely with the identified GEP cell memberships (Figure S3E, S3F), enabling us to confidently assign the two distinct cell-cycle modules specifically to S-phase and G2M-phase, respectively (Figure 3C). The shared CD pathology-associated GEP was active primarily in absorptive lineages in both PDOs and tissues, implicating a conserved cell-type specificity in CD pathogenesis (Figure S3G, S3H). Modules related to intestinal absorption and ribosomal activity were similarly conserved (Figure 3C). However, some GEP modules were specific to the tissue samples, including mucin synthesis and chemosensory function, indicating the absence of secretory lineages in PDO cultures (Figure 3C).

**Figure 3:**
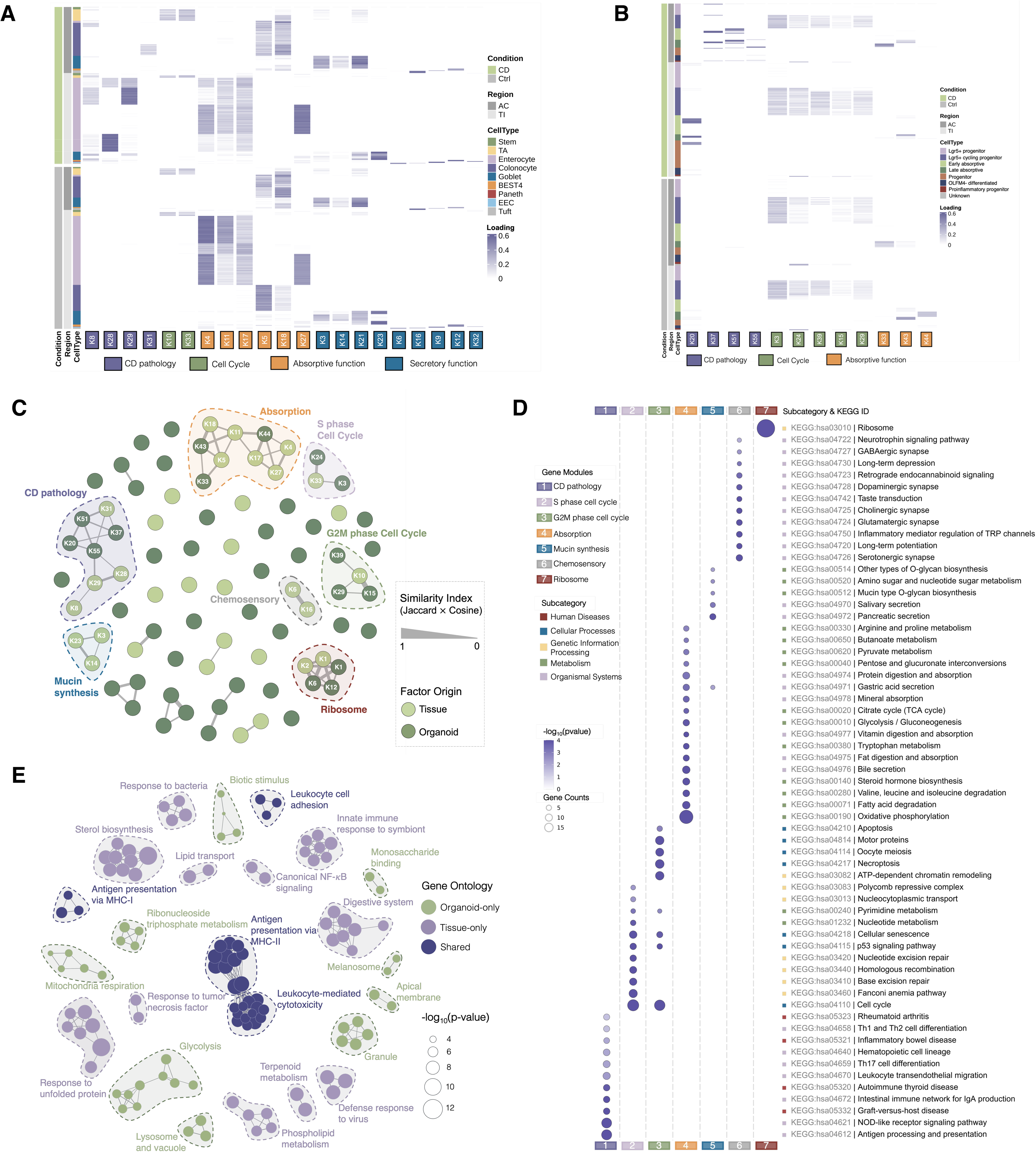
Gene expression programs (GEPs) reveal conserved cell-intrinsic signatures and loss of immune and microbial cross talk in PDOs. (A) Heatmap showing the memberships of tissue-derived cells to GEPs identified from the tissue scRNA-seq data. (B) Heatmap showing the memberships of PDO-derived cells to GEPs identified from the PDO scRNA-seq data. (C) Network clustering of GEPs identified in PDOs and tissues. Each node represents a distinct GEP and is color-coded according to its origin (PDO or tissue). Edges connect GEPs that share driving genes between PDOs and tissues. Edge width corresponds to the similarity index, defined by the Jaccard index of driving gene overlap and cosine similarity between GEPs (only edges with similarity ≥ 0.05 are displayed). (D) KEGG pathway enrichment analysis for selected grouped GEPs (GEP modules). (E) Gene Ontology enrichment analysis highlighting shared and unique driving genes within PDO- and tissue-derived GEP modules associated with CD pathology. Each node in the network represents a Gene Ontology term, and edges are drawn between nodes whose gene sets share more than 50% overlap.

To functionally characterize these GEP modules, we performed KEGG pathway enrichment analysis (Figure 3D). CD pathology GEPs were enriched for pathways including antigen processing and presentation, leukocyte recruitment, T^h^ differentiation, and inflammatory bowel disease, consistent with known disease processes in CD pathology. Cell cycle modules were enriched for nucleotide metabolism, *p53* signaling pathways, and DNA repair, while the absorption-related modules showed enrichment for bile secretion, fatty acid metabolism, and protein digestion and absorption. Tissue-specific modules such as mucin synthesis and chemosensory function showed KEGG enrichment in O-glycan biosynthesis and taste transduction, respectively. Ribosome-related modules were uniformly enriched for translation pathways.

We then investigated which genes and biological functions were shared between PDOs and tissues within the CD pathology-associated GEPs. While the driving genes of this module partially overlapped, some were specific to a particular system. The shared genes were broadly distributed across both PDO and tissue-derived factors and represented a conserved core transcriptional program involved in intrinsic epithelial inflammatory responses (Figure S3I). To examine the biological roles of both shared and system-specific genes, we performed Gene Ontology enrichment analysis (Figure 3E). Shared driving genes were significantly enriched in antigen presentation, leukocyte adhesion, and leukocyte-mediated cytotoxicity pathways, consistent with our previous findings from DEG analysis (Figure 3E, 2G). Tissue-specific driving genes showed enrichment for processes such as defense response to bacteria and virus, innate immune response to symbionts, and tumor necrosis factor signaling, presumably reflecting the influence of immune and microbial interactions present only in the tissue microenvironment. Conversely, driving genes specific to PDOs-cultured in isolation from all such influences-were enriched for pathways related to bioenergetic metabolisms such as monosaccharide binding, glycolysis, and mitochondria respiration, suggesting metabolic adaption to culture condition.

### Extensive divergence of gene regulatory change in core IBD risk genes between PDOs and tissues

To dissect how regulatory control of IBD risk genes differs between PDOs and tissues, we leveraged scATAC-seq data from matched PDO and tissue cell states, analyzing how these systems diverged in terms of CD-specific shifts in expression and accessibility. For each pair of analogous epithelial states (*LGR5*+ progenitor and stem cells, *LGR5*+ cycling progenitor and TA cells, early/late absorptive and enterocyte/colonocyte) ), we first identified DEGs and differentially accessible regions (DARs) between CD and Ctrl conditions, then performed pairwise comparisons to see whether these disease-specific differences were conserved between tissues and PDOs (Figure 4A). We found that, even when originating from the same patients, PDOs and tissues showed markedly different regulatory landscapes. At the transcriptional level, the systems displayed reasonable concordance, with 722, 873, and 659 overlapping DEGs across the three cell-type pairs, respectively (Figure 4A). However, the overlap among differentially accessible regions (DARs) was dramatically smaller. In the *LGR5*+ progenitor-stem pair, we identified 12,477 DARs in PDOs, but only 2 in tissues, and in the absorptive cell pair, 24,207 DARs in PDOs versus 1,069 in tissues, with only 287 overlapping regions (Figure 4A). This disparity suggests that PDOs and tissues achieved similar gene expression output through distinct regulatory mechanisms.

**Figure 4:**
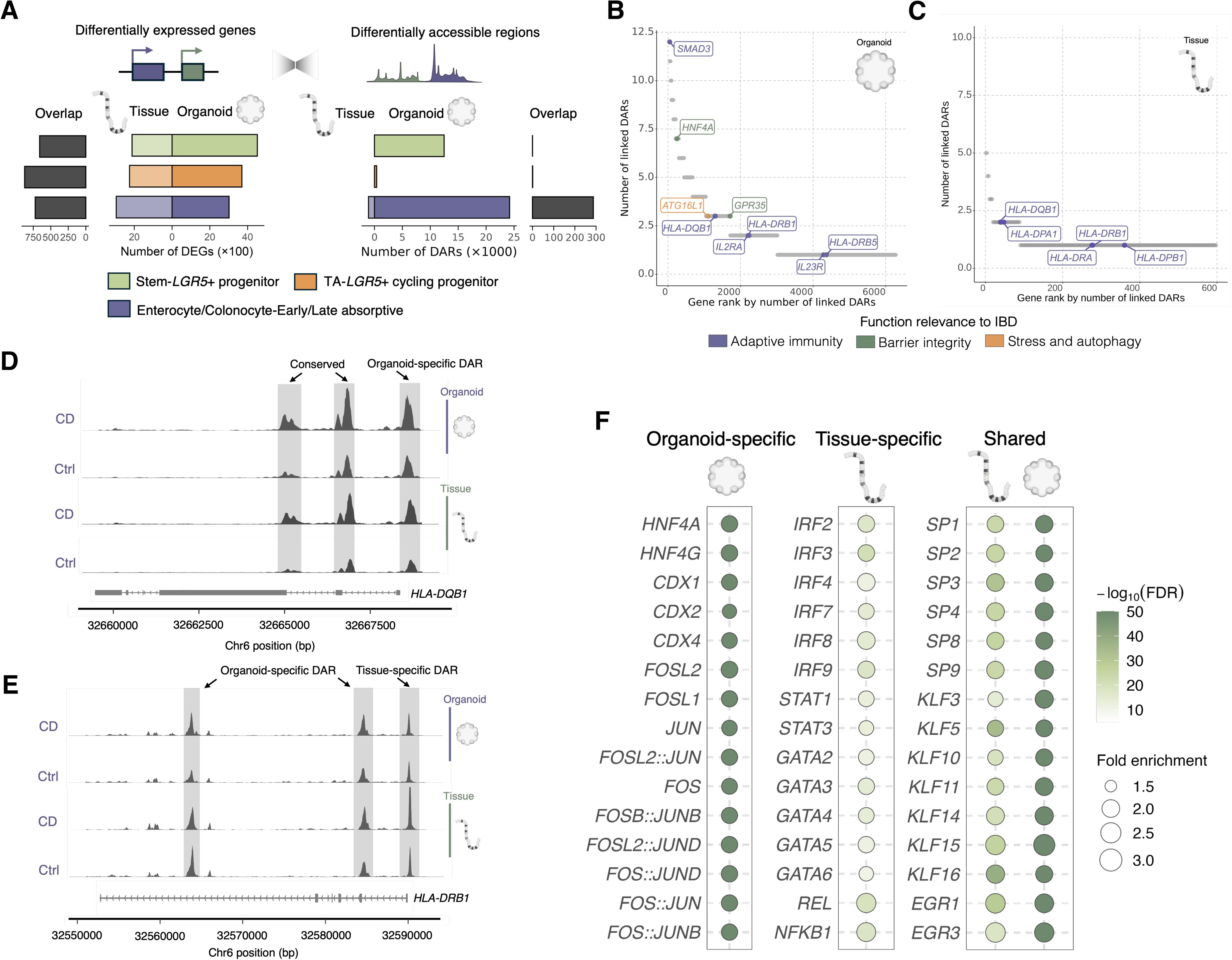
CD-specific chromatin regulatory divergence between PDOs and tissues. (A) Pairwise comparison of the differential expression and differential accessibility across analogous cell states from PDOs and tissues. (B-C) Peak-gene linkage analysis of differential accessible regions (DARs) identified in absorptive cells of PDOs (B) and tissues (C), ranked by number of regulatory regions linked to each gene. (D) A coverage plot showing the chromatin accessibility tracks at the *HLA-DQB1* locus in absorptive cells, showing conserved accessible regions across PDO and tissue. Tracks are separated by source system and disease condition. (E) A coverage plot showing the chromatin accessibility tracks at the *HLA-DRB1* locus in absorptive cells, showing different accessible regions across PDO and tissue. Tracks are separated by source system and disease condition. (F) A dot plot showing the motif enrichment of DARs identified in the absorptive cells from tissue and PDOs. The dot size indicates the fold of enrichment; the color indicates the false discovery rate (FDR).

We performed peak-gene linkage analysis to nominate the putative regulatory targets (Figure 4B-4C), restricting our analysis to absorptive cells since only this group yielded sufficient DARs for meaningful comparisons. As shown in Figure 4D, we identified a tiny set of DARs regulating shared IBD risk gene *HLA-DQB1* (gene body, Chr6: 32664742-32665354 and 32666026-32666898). However, these shared loci represented only a minor fraction of total DARs; the overwhelming majority were system-specific. Notably, even at the *HLA-DRB1* locus, where we observed shared upregulation, two DARs were found to be organoid-specific (Chr6: 32583560-32584808 and 32589425-32590183) and one DAR was found to be tissue-specific (promoter region, Chr6: 32589425-32590183), as shown in Figure 4E. Other PDO-specific DARs (Figure 4B) were linked to adaptive immunity (*IL2RA*, *IL23R*, *SMAD3*), epithelial barrier integrity (*HNF4A, GPR35*), and autophagy (*ATG16L1*), spanning diverse genomic contexts including intronic regions in *HNF4A*, *ATG16L1*, *SMAD3, IL2RA* (Figure S4A-S4D) and promoter-proximal regions of *IL23R* and *SMAD3* (Figure S4C and S4E). In contrast, tissue-specific DARs were associated with antigen presentation genes (*HLA-DRA*, *HLA-DRB1*, *HLA-DPA1*, *HLA-DPB1*), often in promoter-proximal regions (Figure 4E, S4F and S4G). Overall, these examples reflect a broader trend in our dataset, where genes showing PDO-specific upregulation in CD are more frequently associated with PDO-specific DARs, nonetheless tissue-specific upregulation is less commonly linked to tissue-specific DARs. This indicates that chromatin accessibility contributes to context-specific regulation in PDOs but is less predictive of transcriptional differences in tissue, and that PDOs and tissues may achieve similar transcriptional outputs through distinct regulatory architectures.

Motif enrichment analysis reinforced this regulatory dichotomy (Figure 4F). Tissue-specific DARs were enriched for interferon regulatory factor motifs (*IRF2*/*3*/*4*/*7/8*/*9*) and STAT factors (*STAT1*/*3*), implying cytokine/interferon-driven responses arising from tissue-specific immune crosstalk. Conversely, organoid DARs showed enrichment in epithelial development factors *HNF4* and *CDX*, alongside AP-1 family motifs (*JUN* and *FOS*). These factors were also consistently enriched both in absorptive and *LGR5*+ progenitor DARs among PDOs (Figure S4H), reflecting a shared cell-intrinsic regulatory program between multipotent progenitors and differentiated absorptive lineages. Notably, the established role of AP-1 transcription factors in epithelial inflammatory memory^48^ suggests a possible mechanism though which PDOs could retain chromatin states shaped by prior inflammatory exposure even with ablation of the physiological conditions from the tissue context.

### Luminal aspirate stimulation selectively drives disease-associated transcriptional change in CD PDOs

We next sought to understand how PDOs would respond to extrinsic factors-namely, metabolites from intestinal microbiota. These metabolites have been proven to act as signaling bioactive molecules with observable effects on host health.^49,50^ However, finding the appropriate combination of factors for stimulation is challenging as cells *in vivo* are exposed to complex, individualized microbial flora. To approximate this complexity, we elected to expose cells to luminal aspirates collected from a cohort of CD patients under clinical remission (n = 5). This treatment allows us to harness the collective properties of the luminal environment, which are unavailable in reductionist single-metabolite treatment and to account for the combined heterogeneity of inter-individual differences and the diversity produced by an entire microbial ecosystem (Figure 5A).

**Figure 5:**
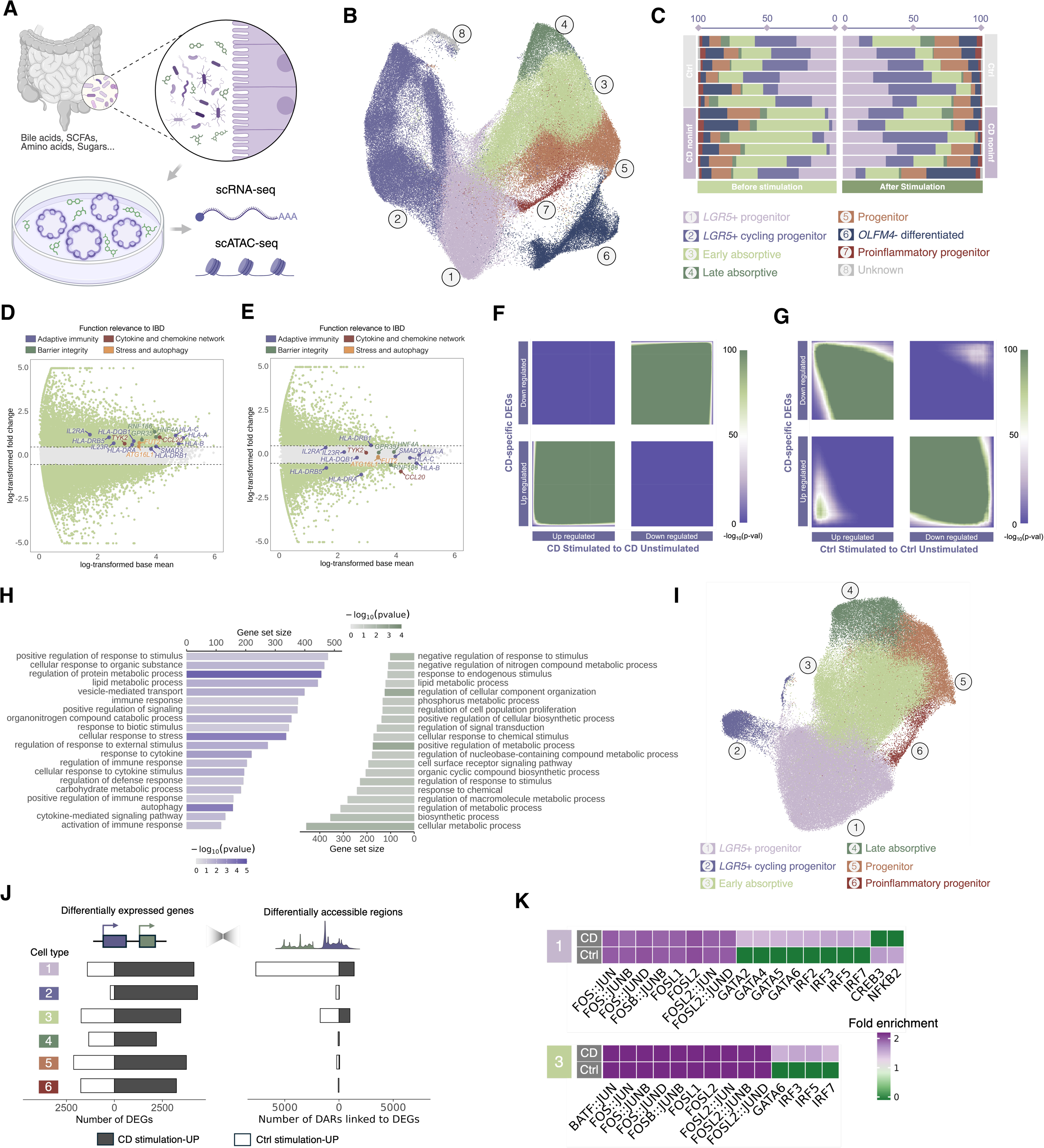
Luminal metabolites selectively activate disease-associated transcriptional and chromatin changes in CD PDOs. (A) Schematic illustration of stimulating PDOs with luminal metabolites and multi-omics profiling (B) UMAP visualization of single-cell RNA sequencing (scRNA-seq) data from stimulated and unstimulated PDOs, colored by identified cell types (C) Cell composition of the individual PDO samples before and after metabolite before and after stimulation. Each bar represents one sample. (D-E) MA plots displaying stimulation-induced differentially expressed genes (DEGs) in (D) CD PDOs and (E) non-IBD control PDOs. (F-G) RRHO2 plots comparing stimulation-induced transcriptional changes in (F) CD PDOs versus CD disease signatures and (G) in non-IBD control PDOs versus CD disease signatures. Heatmap color indicates the significance (–log₁₀ p-value) of gene-list overlap determined by hypergeometric tests. (H) Gene set enrichment analysis comparing transcriptional changes of stimulation in CD (left) and non-IBD control (right) PDOs. (I) UMAP visualization of single-cell ATAC sequencing (scATAC-seq) data from stimulated PDOs and unstimulated controls, colored by identified cell types. (J) Pairwise comparison of the differential expression and differential accessibility across various cell states from PDOs. Only regions linked to upregulated genes are shown. (K) A heatmap showing the motif enrichment of stimulation-induced DARs from *LGR5*+ progenitor and early absorptive cells, separated by disease condition. The color indicates the fold of enrichment.

Luminal aspirates were collected from both colon and ileum for region-matched stimulation to prevent off-target modulatory biases.^51^ We stimulated PDOs derived from non-IBD controls and CD noninf patients with these luminal aspirates to simulate baseline epithelial responses in the absence of active inflammation. Targeted metabolomics recovered more than 50 types of known human-derived or microbe-modified metabolites from a pool of 300+ types of detectable compounds (Figure S5A). These metabolites included bile acids, short-chain fatty acids, indoles, and amino acids. Untargeted metabolomics further suggested the putative inclusion of lipids, peptides, and other bioactive molecules, providing a complex metabolic milieu representative of the intestinal luminal environment.

To characterize the epithelial response to luminal metabolites, we performed scRNA-seq on stimulated PDOs (121,936 cells) and integrated the data with unstimulated controls (74,046 cells). UMAP visualization and cell type annotation revealed consistent marker gene expression as described above (Figure 5B, S5B). We noted a significant increase in proliferation among CD PDOs following stimulation (Figure 5C), as indicated by a greater proportion of regenerative *LGR5*+ and cycling progenitors (Dirichlet regression, p = 3.8 × 10^-3^). Control PDOs, in contrast, showed no significant compositional changes (Dirichlet regression, p = 6.4 × 10^-1^).

Differential expression analysis revealed that many core IBD risk genes were significantly upregulated in CD PDOs after stimulation (Figure 5D), including MHC-I (*HLA-A*, *HLA-B*, *HLA-C*), MHC-II (*HLA-DRA*, *HLA-DRB5*, *HLA-DQB1*), and other genes that are functionally relevant to adaptive immunity (*SMAD3*, *IL23R*, *IL2RA*), barrier integrity (*HNF4A*, *RNF186*, *GPR35*), cytokine and chemokine signaling (*CCL20*, *TYK2*), stress and autophagy (*FUT2*, *ATG16L1*). In contrast, Ctrl PDOs showed minimal changes in these genes following stimulation (Figure 5E). To assess the transcriptional concordance between gene expression in stimulated PDOs and transcriptional signatures of disease, we performed a Pearson correlation and RRHO2 analysis comparing stimulation-induced DEGs against CD-specific signatures obtained by contrasting unstimulated CD to Ctrl PDOs. Pearson correlation captures global similarity in gene expression changes, while RRHO2 evaluates the significance of overlap between ranked gene lists. This analysis demonstrated strong positive concordance between CD PDO stimulation DEGs and CD-specific signatures (ρ=0.62, Figure 5F, S5C), while Ctrl PDO stimulation DEGs showed discordance with disease signatures (ρ=-0.33, Figure 5G, S5D).

Gene set enrichment analysis further revealed divergent pathway responses between CD and Ctrl PDOs. While both exhibited metabolic change as expected from metabolite exposure, CD PDOs were uniquely enriched for IBD-associated pathways, including immune response, cytokine signaling, and stress and autophagy response (Figure 5H). Contrarily, Ctrl PDO responses were predominantly limited to metabolic adaptation pathways (Figure 5H). These findings demonstrated that CD PDOs are primed to mount disease-relevant response to the same extrinsic stimuli, suggesting disease-relevant transcriptional responses are a convergent result from both cell-intrinsic and cell-extrinsic cues.

### CD PDOs exhibit environment-responsive inflammatory chromatin remodeling upon stimulation

We next investigated the underlying regulatory changes driving divergent transcriptional responses between CD and Ctrl PDOs. To do so, we performed scATAC-seq on matched stimulated (90,524 nuclei) and unstimulated PDOs (69,244 nuclei) from above and integrated the datasets. UMAP visualization revealed major cell states largely consistent with our previous scATAC-seq annotations, including *LGR5*+ progenitors, *LGR5*+ cycling progenitors, early/late absorptive, progenitors, and proinflammatory progenitors (Figure 5I). Peak calling was performed using MACS2^52^ stratified by both cell types and conditions (Figure S5E), yielding a union sets of 442,688 peaks. Accessibility across different cell types at marker gene loci recapitulated prior patterns (Figure S5F-S5I). Notably, the increased cell numbers after introducing stimulated samples allowed us to recover the rare proinflammatory progenitor cluster, characterized by accessible chromatin at the *CCL20* locus (Figure S5I), which was not detected in unstimulated PDOs (Figure 1H).

We next computed DEGs and DARs in matched cell types between stimulated and unstimulated conditions in matched cell types. Differential expression analysis demonstrated that CD PDOs were transcriptionally more responsive to stimulation than Ctrl PDOs, exhibiting a greater number of DEGs across multiple cell types (Figure 5J). However, regarding chromatin accessibility changes, we detected significant DARs only in *LGR5*+ progenitor and early absorptive cells; they were absent in more differentiated cell types (Figure 5J). We speculated that although stimulation broadly affects gene expression, chromatin remodeling may be concentrated in progenitor populations with more epigenetic plasticity.

To identify the regulators driving these chromatin changes, we performed motif enrichment analysis on DARs within *LGR5*+ progenitor and early absorptive cells. In *LGR5*+ progenitor from CD, accessible regions were enriched for motifs of environment-responsive transcription factors which were identified in CD tissue epithelium, including *GATA2*/*4*/*5*/*6*, *STAT3*, and *IRF2*/*3*/*5*/*7* (Figure 5K). These factors are known targets of inflammatory cytokine signaling. Similarly, CD absorptive progenitors showed enrichment for *GATA6* and *IRF3*/5/*7* motifs, indicating sustained sensitivity to inflammatory signals along the differentiation trajectory (Figure 5K). In contrast, Ctrl PDOs showed no enrichment in these motifs. *LGR5*+ progenitors in the Ctrl condition were enriched for stress-related motifs, such as *CREB3* and *NFKB2* (Figure 5K), suggesting a response to luminal stimuli that does not engage inflammatory programs. AP-1 family motifs (*FOS* and *JUN*) were enriched across both CD and Ctrl PDOs and in both cell types (Figure 5K), representing a conserved response to stimulation that is independent of disease state. Taken together, these findings outline a persistent CD-specific pattern of responsivity to cell-extrinsic environmental factors that is partially coordinated by chromatin remodeling at inflammation-sensitive genetic loci.

## Discussion

Relapsing inflammation that involves a wide array of cell types is a hallmark of Crohn’s disease, yet the biological basis for its persistence and recurrence remains incompletely understood. Single-cell atlases have revealed stable transcriptional alterations in different cell compartments even during clinical remission, suggesting that disease-relevant cellular programs may be maintained independently of inflammatory triggers accompanying active inflammation. However, transcriptional profiling of *in vivo* samples cannot by itself reveal whether such alterations are solely autonomous or perpetuated by inflammatory and microbial signals present even during remission.

Our work addresses this question by integrating patient-matched organoid models with luminal metabolite stimulation, as well as single-cell transcriptome and chromatin accessibility profiling to dissect the relative contributions of cell-intrinsic and environment-responsive factors to epithelial dysfunction in CD. In organoids isolated from the intestinal environment, we identified disease-linked transcriptomic differences that support a qualitative measure of the disease, implying that these cell-intrinsic features constitute a portion of the overall landscape of the disease activity. Additional disease signatures detected in tissue alone represent the integration of external signals from immune and microbial cells. An interesting observation is that CD PDOs can uncover disease signatures that are originally masked by a complex tissue environment, such as proliferation defects in stem cell compartments, selective upregulation of select IBD risk genes, and altered accessibility at *cis*-regulatory elements controlling these genes (e.g. *IL23R*, *ATG16L1*). This may also suggest the compensatory signals present *in vivo* can partially correct the disease-associated epithelial dysfunction. Tissue and organoid systems also displayed a striking difference in major histocompatibility complex (MHC) activity: while we observed a ubiquitous increase of MHC-II expression in diseased tissues consistent with previous reports,^2,53^ the expression pattern of these genes in CD organoids was heterogeneously sparse. One possible explanation is that the distribution of the adaptive immune ‘memory’ that triggers MHC activity is uneven; another is that the activation of this immune signature is intrinsically stochastic. Finally, we demonstrated that exposure to physiologically relevant luminal metabolites selectively triggers disease-related transcriptional and epigenetic programs in CD but not control organoids. This difference in responsivity offers evidence that the luminal stimuli could direct CD epithelium towards a preferred proinflammatory state that is not activated in non-IBD cells. Collectively, these findings support a model in which persistent epithelial dysfunction in CD is maintained by both intrinsic susceptibility and extrinsic environmental stimulation.

This work has several implications for understanding CD pathogenesis and relapse. First, it suggests that epithelial dysfunction in CD may not be a sole consequence of maladaptive immune responses to external stimuli; rather it may be partially explained by cell-autonomous alterations that persist independently. Second, epithelial organoid response to the luminal triggers is shaped by disease history, raising the possibility that relapse in CD may be partially driven by persistent epithelial hypersensitivity to luminal cues. Third, our findings imply that long-term disease control may require not only immunosuppression agents, but also additional intervention to restore normal epithelial homeostasis and responses to environmental cues.

We envisage a few directions for future work. First, our findings uncovered a set of CD-specific intrinsic features that persist independently of inflammation, presenting the immediate question of whether such alterations are unique to CD, or shared with other chronic inflammatory disorders, such as ulcerative colitis and celiac disease. Ulcerative colitis, despite sharing clinical features with CD, exhibits distinct pathology such as continuous mucosal inflammation restricted to the colon, as well as divergent genetic susceptibility.^54–56^ Comparative multi-omics profiling would reveal whether epithelial alterations in these conditions are disease-specific or involve a common inflammatory memory program. Second, the discovery that CD PDOs are hypersensitive to luminal stimuli raises the question that whether epithelial homeostasis can be therapeutically restored. Current therapies for CD primarily target immune effector mechanisms such as anti-TNF, anti-IL12/23 and anti-integrin, none of which directly address epithelial dysfunction. Future work should aim to map the molecular vulnerabilities within these CD-specific programs in epithelial cells to identify candidate pathways whose inhibition suppresses disease-biased responses while preserving epithelial integrity. Finally, our results suggest that CD epithelial cells retain a form of persistent immunological adaptation that shapes their response to environmental stimuli even when inflammation is resolved. Elucidating the mechanisms underlying this phenomenon will be an important thrust. Previous studies have reported similar behavior in other barrier tissues,^57,58^ such as skin^59^ and respiratory^60^ epithelial cells, however, little is known about their similarities or differences in specificity or durability. Longitudinal organoid culture systems, high-throughput stimulation designs, and single-cell lineage tracing may be crucial in determining how inflammatory exposures are encoded, maintained, and transmitted across epithelial generations. Such insights will be essential for understanding the persistence of chronic inflammatory disorders persist and for developing interventions that erase maladaptive epithelial memory.

### Limitations of the study

Although this study demonstrates the conserved and distinct physiology of organoids and their matched tissues, some limitations leave certain questions unresolved for future studies. First, the organoids were grown in expansion media designed to sustain intestinal stem cells, and thus the cellular composition of the organoids in this study is predominantly proliferating progenitors and stem cells, which do not fully recapitulate the cellular diversity and functional maturity of the human intestinal epithelium *in vivo*. Future studies employing differentiation protocols, such as withdrawal of Wnt ligands and R-spondin to promote enterocyte differentiation, or addition of Notch inhibitors to enrich for secretory lineages, would provide a more comprehensive view of the full diversity of intestinal epithelial cell types. Second, the variability in experimental conditions potentially introduces technical variation that confound biological interpretation. Organoid cultures were maintained using two different expansion media formulations across samples (see *Methods*). Although both media support intestinal organoid growth through activity of Wnt3a, R-spondin, and Noggin signaling, the precise concentrations of growth factors, small molecules, and undefined components may differ between the two formulations. Single-cell RNA sequencing was performed using two different 10x Genomics chemistries: the Next GEM Single Cell 3’ kit and the more recent GEM-X Single Cell 3’ kit. These two platforms differ in their microfluidic chip design and capture efficiency. Thus, samples profiled with different chemistries may exhibit differences in the number of genes and UMIs detected per cel and the representation of lowly expressed transcripts. Such technical differences in library complexity and sensitivity can influence the detection of subtle transcriptional signatures.

## Data availability

The scRNA-seq and scATAC-seq data of organoid were deposited in the Gene Expression Omnibus (GEO) repository under accession code GSE334942. The matched single-cell data of tissue were deposited in the GEO repository under accession code GSE266616 (scRNA) and GSE273194 (scATAC).

## Code availability

Code for data processing and analysis is available on GitHub at https://github.com/CambridgeLiu/Crohns-Disease-Organoid-analysis.

## Acknowledgement

This work is part of the Gut Cell Atlas Crohn’s Disease Consortium and was funded by The Leona M. and Harry B. Helmsley Charitable Trust and is supported by a grant from Helmsley to The University of Chicago (https://helmsleytrust.org/gut-cell-atlas/). The immunofluorescence experiment was supported by the Organoid and Primary Culture Research Core and Human Tissue Resource Center at the University of Chicago. The metabolomics were performed by the Host-Microbe Metabolomics Facility at the University of Chicago. The sequencing was performed by the Genomics Facility at the University of Chicago. Computational analyses were completed using resources provided by the University of Chicago Research Computing Center. We thank Dr. Ran Zhou and Dr. Cambrian Y. Liu for helpful discussion of the analysis, and Dr. Orlando DeLeon for the guidance on stimulation experiments using luminal metabolites.

## Author contributions

J.L. generated organoid data, performed data analysis, and created figures. R.Z. generated primary tissue data. J.K., J.L. and C.M.C. generated organoids. J.L. and A.B. wrote the manuscript. J.K. and C.M.C. contributed to human sample collection. P.C. contributed to data analysis. A.M.S contributed to metabolomics profiling and data analysis. M.S., S.P., E.B.C., and A.B. contributed to data interpretation. A.B. conceptualized and supervised the study.

## Methods

### Human subject

Intestinal biopsy samples were collected during endoscopic screening at the University of Chicago Medical Center. The protocol for tissue collection was approved by the University of Chicago Institutional Review Board (IRB #15573A), and written informed consent was obtained from all participants. Samples, clinical metadata, and demographic data were collected by clinical research coordinators and provided anonymously to the research team. Immediately after tissue collection, de-identified biopsy tissue was placed in ice-cold tissue medium and transported to the Chang laboratory.

Luminal aspirates were collected at the time of endoscopy/colonoscopy, mixed 1:1 with 2× PBS (v/v) (final: 1× PBS). Immediately after collection, luminal aspirates were sterilized through filtration (0.22 μm) and frozen at -80°C.

### Patient-derived organoid culture

Patient-derived organoids were generated as previously described^17^ with minor adaptations. Briefly, ileal or colonic biopsies were rinsed repeatedly in ice-cold PBS and then resuspended in 10 mL ice-cold 8 mM calcium-chelating EDTA in PBS. The tissues were agitated on a rocker at 4 °C for 30 min and replaced in ice-cold Advanced DMEM/F-12 (Life Technologies, cat. #11320033) with gentle pipetting to detach epithelium crypts. The epithelium crypts were filtered (100 µm cell strainer), centrifuged (400 × g, 5 min, 4 °C) and resuspended in a 1:1 mixture of human organoid growth medium^17^ or IntestiCult^TM^ organoid growth medium and Matrigel (growth factor reduced, Corning, cat. #CLS356231). Matrigel domes were plated in 6-well plates and incubated at 37 °C, 5% CO₂ for ∼1 h for gelation, after which organoid culture medium was added. Cultures were maintained with medium changes twice per week and passaged approximately every 2 weeks. The organoid metadata of culture medium, passage number, and processing information is provided in Supplementary Table S1.

For stimulation, luminal aspirates from five CD patients under clinical remission were thawed, averagely mixed, and added to organoid culture medium for a final concentration of 10% (v/v). Organoids were stimulated for 24 hrs.

### Organoid dissociation and nuclei isolation

Matrigel domes were collected and centrifuged at 400 × g for 5 min at 4°C. Media and Matrigel were removed and the cell pellet was rinsed repeatedly in ice-code PBS. Organoids were suspended in 3mL of pre-warmed TrypLE express (Thermo Fisher cat. #12605010) and rotated at 37°C for 10 minutes. Afterwards, organoid cell suspension was triturated with a P1000 pipette. Cells were filtered through a 40 μm cell strainer (Falcon). The cells were then processed using the EasySep™ Dead Cell Removal (Annexin V) Kit following manufacturer instructions to select high-quality viable cells. Final viable cells were resuspended in single-cell suspension buffer recommended by 10x Genomics (PBS and 0.04% BSA) and assessed for viability with Trypan Blue.

Cell nuclei extraction was performed following CG000169 Rev E (10x Genomics). Briefly, 100 μL of lysis buffer was added to suspend cell pellets with gentle pipetting. After 4-min incubation on ice, lysis was neutralized by wash buffer. The nuclei were collected by centrifugation (500 × g, 5 min, 4 °C) and suspended in nuclei buffer. The final concentration of nuclei was adjusted for targeted recovery number.

### Harvesting and Immunofluorescence staining of tissue biopsy and organoids

Tissue biopsies were rinsed repeatedly in ice-cold PBS, fixed using 10% neutral buffered formalin (NBF) for 24 hrs, dehydrated using 70% ethanol, and embedded in paraffin. To harvest organoids cultured in Matrigel domes, cultures were washed with ice-cold PBS and incubated with 2mL of ice-cold Organoid Harvest Solution (Bio-techne cat. #3700-100-01) for 30 min. After the organoid were detached from Matrigel and settled down, they were collected by centrifugation (500 × g, 7 min, 4 °C) and fixed using 10% NBF for 1hr. Fixed organoids were collected by centrifugation (1000 × g, 7 min, 4 °C), stabilized in HistoGel, dehydrated using 70% ethanol, and embedded in paraffin. The embedded specimen blocks were sectioned into 5μm thickness slides.

To perform immunofluorescence staining, the samples were first deparaffinized using HistoClear (National Diagnostics, cat. #50-899-90147) (3 × 5 min), then rehydrated with decreasing concentrations of ethanol, as follows: 100% ethanol (2 × 5 min), 70% ethanol (2 × 5 min), 50% ethanol (2 × 5 min), water (2 × 5 min). Antigen retrieval was performed for 10-15 min in a benchtop cooker with 1× Citrate Buffer, pH 6.0 (Sigma Aldrich, cat. #C9999). Primary antibodies used were anti-OLFM4 (Cell Signaling Technology, cat. #14369), anti-KI67 (Invitrogen, cat. #740008T), anti-ANPEP (Proteintech, cat. #66211-1-Ig), anti-HLA-DRA (Abcam, cat. #ab20181). The staining was imaged using Leica Stellaris 8 Laser Scanning Confocal at the University of Chicago Integrated Light Microscopy Core.

### Single-cell RNA library preparation and sequencing

Single-cell transcriptomic libraries were processed using the Chromium Single Cell 3′ Gene Expression kit v.3.1 or GEM-X Single Cell 3’ Kit v4 (10x Genomics) following manufacturer’s instructions. The libraries were sequenced on Illumina’s NovaSeq X at the Genomics Core Facility at the University of Chicago with a sequencing depth of 50,000 read/cell. For detailed information on each library and associated metadata, refer to Supplementary Table S1.

### Single-cell ATAC library preparation and sequencing

Isolated nuclei were suspended in nuclei buffer at a concentration of 2,500 nuclei/μL. Libraries were prepared using the Chromium Next GEM Single Cell ATAC v2 kit (10x Genomics) following manufacturer’s instructions. The libraries were sequenced on Illumina’s NovaSeq X at the Genomics Core Facility at the University of Chicago with a sequencing depth of 50,000 read/cell.

### Metabolomics of luminal metabolites

#### Extraction

Extraction solvent (80% methanol spiked with internal standards) was added at a ratio of 100 mg of material/mL of extraction solvent in beadruptor tubes (Fisher; cat. #15-340-154). Samples were homogenized at 4 °C on a Bead Mill 24 Homogenizer (Fisher; cat. #15-340-163), set at 1.6 m/s with 6 thirty-second cycles, 5 seconds off per cycle. Samples were then centrifuged at -10 °C, 20,000 × g for 15 min and the supernatant was used for subsequent metabolomic analysis.

#### Short chain fatty acids

Short chain fatty acids were first derivatized as described by Haak *et al*.^61^ with the following modifications. 100 µL metabolite extract was added to 100 µL of 100 mM borate buffer (pH 10) (Thermo Fisher, cat. #28341), 400 µL of 100 mM pentafluorobenzyl bromide (Millipore Sigma; cat. #90257) in Acetonitrile (Fisher; cat. #A955-4), and 400 µL of n-hexane (Acros Organics; cat. #160780010) in a capped mass spec autosampler vial (Microliter; cat. #09-1200). Samples were heated in a thermomixer C (Eppendorf) to 65 °C for 1 hour while shaking at 1300 rpm. After cooling to RT, samples were centrifuged at 4°C, 2,000 × g for 5 min, allowing phase separation. The hexanes phase (100 µL, top layer) was transferred to an autosampler vial containing a glass insert and the vial was sealed. Another 100 µL of the hexanes phase was diluted with 900 µL of n-hexane in an autosampler vial. Concentrated and dilute samples were analyzed using a GC-MS (Agilent 7890A GC system, Agilent 5975C MS detector) operating in negative chemical ionization mode, using a HP-5MSUI column (30 m × 0.25 mm, 0.25 µm; Agilent Technologies 19091S-433UI), methane as the reagent gas (99.999% pure) and 1 µL split injection (1:10 split ratio). Oven ramp parameters: 1 min hold at 60 °C, 25 °C per min up to 300 °C with a 2.5 min hold at 300 °C. Inlet temperature was 280 °C and transfer line was 310 °C. A 10-point calibration curve was prepared with acetate (100 mM), propionate (25 mM), butyrate (12.5 mM), and succinate (50 mM), with 9 subsequent 2× serial dilutions. Data analysis was performed using MassHunter Quantitative Analysis software (version B.10, Agilent Technologies) and confirmed by comparison to authentic standards. Normalized peak areas were calculated by dividing raw peak areas of targeted analytes by averaged raw peak areas of internal standards.

#### Bile acids

Bile acids were analyzed using LC-MS.^62^ 75 µL metabolite extract was added to prelabeled mass spectrometry autosampler vials (Microliter; cat. #09-1200) and dried down completely under a nitrogen stream at 30 L/min (top) 1 L/min (bottom) at 30 °C (Biotage SPE Dry 96 Dual; 3579M). Samples were resuspended in 50:50 Water:Methanol (750 µL). Vials were added to a thermomixer C (Eppendorf) to resuspend analytes at 4 °C, 1,000 rpm for 15 min with an infinite hold at 4 °C. Samples were then transferred to prelabeled microcentrifuge tubes and centrifuged at 4 °C, 20,000 × g for 15 min to remove insoluble debris. The supernatant (700 µL) was transferred to a fresh, prelabeled mass spectrometry autosampler vial. Samples were analyzed on a liquid chromatography system (Agilent 1290 infinity II) coupled to a quadrupole time-of-flight (QTOF) mass spectrometer (Agilent 6546), operating in negative mode, equipped with an Agilent Jet Stream Electrospray Ionization source. The sample (5 µL) was injected onto an XBridge© BEH C18 Column (3.5 µm, 2.1 × 100 mm; Waters Corporation, PN) fitted with an XBridge© BEH C18 guard (Waters Corporation, PN) at 45 °C. Elution started with 72% A (Water, 0.1% formic acid) and 28% B (Acetone, 0.1% formic acid) with a flow rate of 0.4 mL/min for 1 min and linearly increased to 33% B over 5 min, then linearly increased to 65% B over 14 min. Then the flow rate was increased to 0.6 mL/min and B was increased to 98% over 0.5 min and these conditions were held constant for 3.5 min. Finally, re-equilibration at a flow rate of 0.4 mL/min of 28% B was performed for 3 min. The electrospray ionization conditions were set with the capillary voltage at 3.5 kV, nozzle voltage at 2 kV, and detection window set to 100-1700 m/z with continuous infusion of a reference mass (Agilent ESI TOF Biopolymer Analysis Reference Mix) for mass calibration. A ten-point calibration curve was used for quantitation. Data analysis was performed using MassHunter Profinder Analysis software (version B.10, Agilent Technologies) and confirmed by comparison with authentic standards. Normalized peak areas were calculated by dividing raw peak areas of targeted analytes by averaged raw peak areas of internal standards.

#### Indoles

Indole-containing metabolites, B-vitamins and other targeted metabolites were analyzed by LC-MS/MS.^63^ 400 µL metabolite extract was added to pre-labeled microcentrifuge tubes. Samples were dried down completely using a Genevac EZ-2 Elite. Samples were resuspended in 100 µL of 50:50 Water:Methanol and added to an Eppendorf thermomixer® C at 4 °C, 1000 rpm for 15 min to resuspend analytes. Samples were then centrifuged at 4 °C, 20,000 × g for 15 min to remove insoluble debris. The supernatant (80 µL) was transferred to a fresh, prelabeled MS vial with inserts or 96 deep-well plate (Agilent 5065-4402). Samples were analyzed on an Agilent 1290 infinity II liquid chromatography system coupled to an Agilent 6470 triple quadrupole mass spectrometer, operating in positive mode, equipped with an Agilent Jet Stream Electrospray Ionization source. Each sample (2 µL) was injected into a Acquity UPLC HSS PFP column, 1.8 µm, 2.1 × 100 mm (Waters; 186005967) equipped with a Acquity UPLC HSS PFP VanGuard Pre-column, 100Å, 1.8 μm, 2.1 mm × 5 mm (Waters; 186005974) at 45 °C. Mobile phase A was 0.35% formic acid in Water and mobile phase B was 0.35% formic acid in 95:5 Acetonitrile:Water. The flow rate was set to 0.5 mL/min starting at 0% B held constant for 3 min, then linearly increased to 50% over 5 min, then linearly increased to 95% B over 1 min, and held at 100% B for the next 3 min. Mobile phase B was then brought back down to 0% over 0.5 min and held at 0% for re-equilibration for 2.5 min. The QQQ electrospray conditions were set with capillary voltage at 4 kV, nozzle voltage at 500 V, and Dynamic MRM was used with cycle time of 500 ms. Transitions were monitored in positive mode for 46 analytes (table on next slide). An 11-point calibration curve (ranging from 0.88 nM to 909 µM) was prepared for tryptophan, tyrosine, phenylalanine, serotonin, 5-HIAA, melatonin, tryptamine, kynurenine, kynurenic acid, anthranilic acid, and niacin. Data analysis was performed using MassHunter Quant software (version B.10, Agilent Technologies) and confirmed by comparison with authentic standards. Normalized peak areas were calculated by dividing raw peak areas of targeted analytes by averaged raw peak areas of internal standards.

#### Untargeted metabolomics

Metabolites were analyzed using GC-MS with electron impact ionization.^64^ 100 µL metabolite extract in mass spec autosampler vials (Microliter; 09-1200) was dried down completely under nitrogen stream at 30 L/min (top) 1 L/min (bottom) at 30 °C (Biotage SPE Dry 96 Dual; 3579M). Dried samples were augmented with 50 µL of freshly prepared 20 mg/mL methoxyamine (Sigma; 226904) in pyridine (Sigma; 270970) was added and incubated in a thermomixer C (Eppendorf) for 90 min at 30 °C and 1400 rpm. After samples are cooled to room temperature, 80 µL of derivatizing reagent (BSTFA + 1% TMCS; Sigma; B-023) and 70 µL of ethyl acetate (Sigma; 439169) were added and samples were incubated in a thermomixer at 70 °C for 1 hour and 1400 rpm. Samples were cooled to RT and 400 µL of Ethyl Acetate was added to dilute samples. Turbid samples were transferred to microcentrifuge tubes and centrifuged at 4 °C, 20,000 × g for 15 min. Supernatants were then added to mass spec vials for GCMS analysis. Samples were analyzed using a GC-MS (Agilent 7890A GC system, Agilent 5975C MS detector) operating in electron impact ionization mode, using a HP-5MSUI column (30 m x 0.25 mm, 0.25 µm; Agilent Technologies 19091S-433UI) and 1 µL injection. Oven ramp parameters: 1 min hold at 60 °C, 16 °C per min up to 300 °C with a 7 min hold at 300 °C. Inlet temperature was 280 °C and transfer line was 300 °C. Data analysis was performed using MassHunter Quantitative Analysis software (version B.10, Agilent Technologies) and confirmed by comparison to authentic standards. Normalized peak areas were calculated by dividing raw peak areas of targeted analytes by averaged raw peak areas of internal standards. A comprehensive list with detected metabolites using untargeted metabolomics is provided in Supplementary Table S6.

### Single cell RNA-seq data pre-processing

scRNA-seq FASTQ files were aligned to the GRCh38 human reference genome to generate the cell-by-gene count matrices, following the Cell Ranger pipeline (10x Genomics). All cell-by-gene count matrices were processed using Scrublet doublet detection pipeline^65^ with a threshold of 0.25, and predicted doublets were removed prior to further analysis. Downstream analysis was conducted using the Seurat R package (v5.1.0), following a pipeline of quality control, data normalization, dimension reduction, data integration, clustering, and cell annotation. We selected high-quality cells with gene features between 200 and 10,000, unique molecular identifiers (UMIs) between 1,500 and 50,000, and mitochondrial gene count percentage less than 25%. Raw count matrices were then normalized by total UMI number per cell, converted to transcripts-per-10,000, and log-transformed. After normalization, top 3,000 highly variable genes were selected by variance stabilizing transformation. The log-normalized data were scaled and subjected to principal component analysis (PCA) using the variable genes. The top 30 PCs were used for downstream analysis. To account for batch effects and harmonize samples across different conditions, we performed data integration using Seurat’s reciprocal PCA (RPCA) implementation.^30^ After data integration, cell clustering was computed using the Louvain algorithm in the RPCA space on the top 30 PCs, followed by FindClusters with resolution = 0.8. Cell embeddings were visualized by Uniform Manifold Approximation and Projection (UMAP) on the top 30 PCs.

### Single cell ATAC-seq data pre-processing

scATAC-seq FASTQ files were aligned to the GRCh38 human reference genome to generate the cell-by-gene count matrices, following the Cell Ranger ATAC pipeline (10x Genomics). The cell-by-peak count matrices were processed using the Signac (v1.16.0) pipeline. To select high-quality nuclei, we kept nuclei with fragments between 1,500 to 15,000, TSS enrichment score greater than 3, and percentage of reads-in-peaks greater than 40%. Raw count matrices were then normalized and dimensionally reduced by latent semantic indexing (LSI, combined steps of TF-IDF followed by singular value decomposition). The first LSI component was discarded as it correlated strongly with sequencing depth; therefore, components 2-30 were used for downstream analysis. To account for batch effects and harmonize samples across different conditions, we performed data integration using Signac’s reciprocal LSI implementation. After data integration, cell clustering was computed using the Louvain algorithm with resolution = 0.6. Cell embeddings were visualized by UMAP on the 2-30 components.

Annotation of PDO nuclei was performed according to a method introduced by Uzquiano *et al*.^66^ Briefly, gene activity scores were computed using Signac’s GeneActivity() function and log-normalized. First, the gene activity data were integrated with the corresponding scRNA-seq into low-dimensional CCA space, then the TransferData() function was used to predict cell type labels for the ATAC cells. We then separately called the differentially accessible regions (DARs) for each scATAC-seq cluster using FindMarkers() with the Wilcoxon rank-sum test. These DARs were mapped to the closest genes, and the top-ranked genes per cluster were used to confirm and refine cell type assignments derived from label transfer.

### SCENT signaling entropy quantification

Signaling entropy, a proxy for cellular differentiation potential, was quantified for epithelial cells derived from both organoid and tissue samples using the SCENT^26,27^ R package (v 1.0.3). Scores were computed using SCENT’s correlation-based approximation, which integrates transcriptomic data with a curated protein-protein interaction network (net17Jan16, human version) derived from Pathway Commons on the 17th of January 2016. Correlation coefficient between signaling entropy scores and crypt-villus axis scores (see below) of organoids were compared using Pearson correlation implemented in the stat_cor() function from the *ggpubr* package.

### Crypt-villus axis score

To quantify the differentiation state of epithelial cells along the crypt-villus axis, we calculated an axis score based on the expression of previously defined marker genes obtained from established crypt-villus zonation studies^28,29^. The gene set included: *SEPP1, CEACAM7, PLAC8, CEACAM1, TSPAN1, CEACAM5, CEACAM6, IFI27, DHRS9, KRT20, RHOC, CD177, PKIB, HPGD, LYPD8, APOBEC1, APOB, APOA4, APOA1, NPC1L1, EGFR, KLF4, ENPP3, NT5E, SLC28A2, ADA*. The expression of these markers was aggregated to generate a continuous score reflecting each cell’s position along the crypt-villus axis, with higher values indicating greater villus-like differentiation.

### Dirichlet regression

Compositional constraint is a major analytical challenge in comparing cell type proportions from single-cell data. The cell proportions within each sample must sum to one, rendering them mutually dependent. We addressed this challenge using a Dirichlet-multinomial regression model, which accounts for the multivariate and compositional nature of the data. This approach has been previously validated for analyzing cell composition changes across conditions.^2,33^

To enable a cross-source comparison between organoids and primary tissue, we grouped epithelial cell types into three biologically meaningful categories: (i) progenitors, which included *LGR5*+ progenitor and cycling progenitor in organoid, and stem and TA in tissue, (ii) absorptive cells, which included early absorptive and late absorptive cells in organoid, and enterocyte and colonocyte in tissue, and (iii) other cells, encompassing all remaining epithelial types. The regression was performed using R package *DirichletReg* (v 0.7-2).

### Differentially expressed gene analysis

To perform differential gene expression (DE) analysis, we first pseudobulked gene counts at the sample level by summing together gene counts of all the cells from the same sample. This approach collapses single-cell data into one aggregated expression vector per sample. We then performed DE analysis using DESeq2^67^. The DE results comparing tissue and PDO are summarized in Supplementary Table S3.

### Rank-rank hypergeometric overlap

To assess the concordance between transcriptional changes from two parallel comparisons, we performed a Rank-Rank Hypergeometric Overlap (RRHO2) analysis^39,40^. This method quantifies the extent of concordance between two ranked gene lists and is particularly well-suited for comparing genome-wide differential expression signatures without relying on selected thresholds.

DEG signatures were identified for matched epithelial cell populations in organoid and tissue datasets using the Seurat FindMarkers() function with default parameters and then ranked by degree of differentiation [-log_10_(p-value) ∗ sign(effect)]. The RRHO2 analysis was then applied to the two ranked lists using a sliding window (step size = 1) across all genes to detect regions of statistically significant overlap. This approach highlights gene expression programs that are consistently upregulated or downregulated in both systems.

### Disease score

The inflammation scoring method is based on the gene signature defined in the TAURUS study^5^, the biopsy-based biomarker of inflammation study^37^, the genome-wide association study (GWAS)^38^ of autoimmune disease, and our previous single-cell atlas study of Crohn’s disease^3^. Briefly, single-cell gene expression data were pseudobulked at the sample level by summing gene counts across all cells within each sample, generating a single expression profile per sample. We then used the refined disease gene set as gene signatures and applied the escape.matrix() function from the *escape* package^68^ with ssGSEA implementation^69^. The scores were scaled between 0 to 10, resulting in a vector representing enrichment of the inflammation score per sample.

### Non-negative matrix factorization of gene expression data

The raw count matrix was normalized by library size to obtain shifted log counts according to Ahlmann-Eltze *et al*.^70^, using the formula

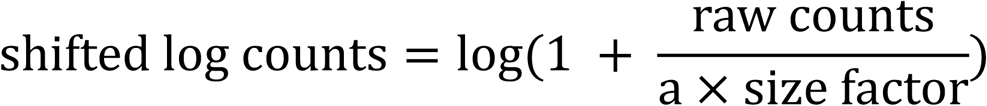

where size factor was calculated as the row sums of the count matrix (library size for each cell) divided by the mean row sum, and a=1 is a pseudo-count parameter.

We applied non-negative matrix factorization (NMF) to the shifted log counts using the *flashier* package (version 1.0.55).^42–44^ This approximated the n × m matrix of shifted log counts, X, where n is the number of cells and m is the number of genes, by a linear combination of K factors, X ≈ LF^T^, where L is the n × K “cell matrix” containing memberships of cells to GEPs, and F is the m × K “gene matrix” containing log-fold expression increases in each GEP. Briefly, we fit the NMF by performing an initial greedy factorization using empirical Bayes matrix factorization with point-exponential priors, which enforce non-negativity constraint. The greedy search extracted up to 100 factors without null checking or backfitting using a homoscedastic variance model. The model was refined through two phases of iterative backfitting, which consisted of an initial set of unaccelerated alternating updates of loadings, and a second set of updates that used the extrapolation method described by Ang *et al*^71^ to accelerate the updates.

For interpreting the NMF results, we note that the measurement scale for each factor k cannot be estimated from the data. This means that, for a given factor k, the memberships and LFCs are only known in relative terms. This also means that memberships and LFCs cannot be compared across factors. Our convention is to rescale the memberships, separately for each factor k, so that the largest (relative) membership is always 1.

To identify the driving genes for each gene expression program (GEP), we leveraged the factor matrix F returned from above computation. Two complementary measures were used to nominate the driving genes: genes with highest values (i.e., largest increases in expression), as well as genes with the largest values relative to all other GEPs (most “distinctive” changes in expression). For each program, we selected the top 100 genes with largest effect sizes and the top 100 most distinctive genes and used the union as the driving gene set representing that GEP.

We quantified pairwise similarity among GEPs obtained from PDO and tissue gene expression data using two complementary metrics: one, Jaccard Index (degree of overlap) in driving genes; two, cosine similarity between the respective columns of the two gene matrices (F). GEPs with combined similarity ≥ 0.05 were considered related and visualized as edges in network analyses (Figure 3C).

The GEPs associated with CD pathology were identified by high membership enrichment in samples derived from CD patients and significant overlap of driving genes with known GWAS-identified CD risk loci. To quantify this, GEP membership scores obtained from the membership matrix L were binarized to select cell that were active for each GEP using a threshold 0.1 of membership scores. We then performed a binomial test for each GEP to assess enrichment of active cells in CD samples leveraging sample metadata. Meanwhile, we assessed each GEP’s driving genes overlap with established IBD genetic risk loci by Fisher’s exact test. GEPs demonstrated both enrichments were classified as CD-associated.

Annotation of GEP modules was performed using Kyoto Encyclopedia of Genes and Genomes (KEGG) with R package *clusterProfiler* (v 4.10.1). Enrichment was assessed with parameters minSize=10, maxSize=500, pvalueCutoff = 0.05. Pathways with FDR < 0.05 were considered significantly enriched. To study the difference of CD GEPs between organoid and tissue, we compared the driving gene compositions from both and classified driving genes as shared, organoid-specific, and tissue-specific. We performed Gene Ontology (GO) analysis of these categories with R package *clusterProfiler* (v 4.10.1), using parameters minSize=10, maxSize=500, pvalueCutoff = 0.05. Pathways with FDR < 0.05 were considered significantly enriched. The summary of KEGG and GO results were provided in Supplementary Table S4.

### Gene set enrichment analysis of stimulated organoids

Gene set enrichment analysis (GSEA)^72^ was performed on stimulation DEGs using R package *clusterProfiler* (v 4.10.1), with parameters minSize=3, maxSize=500, pvalueCutoff = 0.05, pAdjustMethod = "BH”. The enriched pathways for stimulated CD and control organoids are provided in Supplementary Table S7.

### Candidate TF regulator identification

We used chromVAR implemented in Signac to perform motif enrichment analysis and calculate Z score across all the cells. To better identify functional transcription factors (TFs) in each cell type, we calculated the Pearson correlation between the motif enrichment Z score and scRNA-measured gene expression across the major cell types. The TFs that passed a filtering threshold (Pearson correlation coefficient>0.8) are defined as the candidate TF regulators.

### Differentially accessible region analysis

DARs were called for each comparison using Signac FindMarkers() function with the Wilcoxon rank-sum test. The DARs that passed a filtering threshold (log2 fold change ≥ 0.5 and p.adjust < 0.01) were retained.

### Peak-gene linkage

To identify putative *cis*-regulatory relationships between accessible chromatin regions and gene expression, we performed peak-to-gene linkage analysis using Signac’s Pearson correlation-based workflow. Peak accessibility from scATAC-seq data was correlated with gene activity data across cells, and significant links were identified based on correlation strength and distance constraints (maximum distance 500 kb, correlation threshold > 0.05, FDR < 0.05). The DARs and their putative gene targets are summarized in Supplementary Table S5.

### Motif enrichments for differentially accessible regions

We performed motif enrichment analysis using Signac with the JASPAR2020 *Homo Sapiens* database^73^. Motif occurrences were measured in DARs and background regions using position weight matrices from database, and enrichment was computed using hypergeometric test with Benjamini-Hochberg correction. Significantly enriched motifs were filtered with the following threshold: observed in more than 10% of tested DARs, fold-enrichment > 1.5, and adjusted p.value < 0.01.

## Supplementary Figure Captions

**Figure S1: Multi-omics’ profile of matched tissues, relate to** Figure 1.

(A) UMAP visualization of tissue epithelium, colored by cell types

(B) Marker genes expression of different cell types in tissue

(C) Signaling entropy score of tissue

(D) Crypt-villus score of tissue

(E) Spearman correlation of signaling entropy score and crypt-villus score in PDOs

(F) UMAP visualization of scATAC-seq of tissue epithelium, colored by cell types

(G) EPCAM gene activity inferred from chromatin accessibility

(H-J) Patient-wise variability of scATAC-seq of PDOs, colored by (H) patient ID, (I) disease condition,

(J) sampling region.

(K) Batch-corrected patient-wise variability of scATAC-seq of PDOs

**Figure S2: Comparison of CD-associated changes in PDOs and matched tissues, related to Figure 2**.

(A–B) Bar plots showing the cell-type composition across individual PDO samples profiled by (A) scRNA-seq and (B) scATAC-seq.

(C) A ternary plot illustrating cell-type composition across matched tissue samples, with each datapoint representing an individual sample.

(D) Dirichlet-multinomial regression analysis assessing cell-type composition changes specific to CD tissues.

(E–F) MA plots highlighting CD-specific DEGs identified in PDOs for comparisons between (E) CD adj vs. non-IBD control, and (F) CD noninf vs. non-IBD control.

(G–H) MA plots highlighting CD-specific DEGs identified in matched tissues for comparisons between

(H) CD adj vs. non-IBD control, and (I) CD noninf vs. non-IBD control.

(I-J) Feature plots showing the gene expression of HLA-DRA in a matched tissue and PDO sample. immunofluorescence staining samples HA56

(K) Scatter plot comparing DEGs identified in PDOs versus matched tissues.

(L) Inflammatory disease scoring of tissue samples calculated using curated inflammatory gene signatures. (M-P) RRHO2 plots comparing transcriptional signatures of matched cell types between PDOs and tissues, using DEGs derived from the CD noninf vs. non-IBD control comparison. Heatmap colors represent the significance (–log₁₀ p-value) of overlap between ranked gene lists.

(Q–T) RRHO2 plots comparing transcriptional signatures of matched cell types between PDOs and tissues, using DEGs derived from the CD adj vs. non-IBD control comparison. Color intensity indicates significance (–log₁₀ p-value) of overlap in ranked gene lists.

**Figure S3: Gene expression programs (GEPs) of PDOs and matched tissues (related to Figure 3).**

(A–B) Heatmaps showing the top driving genes and their corresponding effect sizes for each identified GEP in (A) tissues and (B) PDOs.

(C–D) Identification of CD-relevant GEPs in (C) PDOs and (D) tissues. The x-axis indicates the log-transformed p-values of membership enrichment within CD-specific cells (Binomial test), and the y-axis represents the log-transformed p-values of driving gene enrichment among GWAS-identified risk genes (Fisher’s exact test).

(E–F) Visualization of the cell-cycle-associated GEPs in (E) tissues and (F) PDOs. Cell-cycle phases were predicted using Seurat, and cell memberships were assigned based on identified cell cycling GEPs.

(G–H) Cell memberships involved in CD pathology-associated GEPs in (G) PDOs and (H) tissues.

(I) The overlapped driving genes of CD pathology GEPs and their effect sizes from PDOs and tissues.

**Figure S4: CD-specific chromatin regulatory divergence between PDOs and tissues (related to Figure 4).**

(A-G) A coverage plot showing the chromatin accessibility tracks at the core IBD risk gene loci in absorptive cells, showing conserved accessible regions across PDO and tissue. Tracks are separated by source system and disease condition. (A) *HNF4A*, (B) *ATG16L1*, (C) *SMAD3*, (D) *IL2RA*, (E) *IL23R*, (F) *HLA-DPB1*, (G) *HLA-DRA*.

(H) A dot plot showing the motif enrichment of DARs shared between the *LGR5*+ progenitor and absorptive cells from PDOs. The dot size indicates the fold of enrichment; the color indicates the false discovery rate (FDR).

**Figure S5: Multi-omics’ profile of stimulated PDOs with luminal metabolites (related to Figure 5).**

(A) Targeted metabolomics of luminal metabolites isolated from ascending colon and terminal ileum.

(B) A dot plot showing expression levels of selected marker genes across stimulated PDO cell types

(C-D) Scatter plots comparing stimulation-induced transcriptional changes in (C) CD PDOs versus CD disease signatures and (D) in non-IBD control PDOs versus CD disease signatures.

(E) Distribution and quantification of accessible chromatin peaks identified within each cell type and disease condition, categorized by genomic region.

(F-I) Coverage plots illustrating normalized chromatin accessibility at genomic peaks associated with marker genes for each cell type.

