## Supplementary figures and images for "Epithelial-Intrinsic Alterations and Maladaptation to Luminal Metabolites Underlie Persistent Crohn’s Disease Pathogenesis"

### Supplementary Figure S1-S5

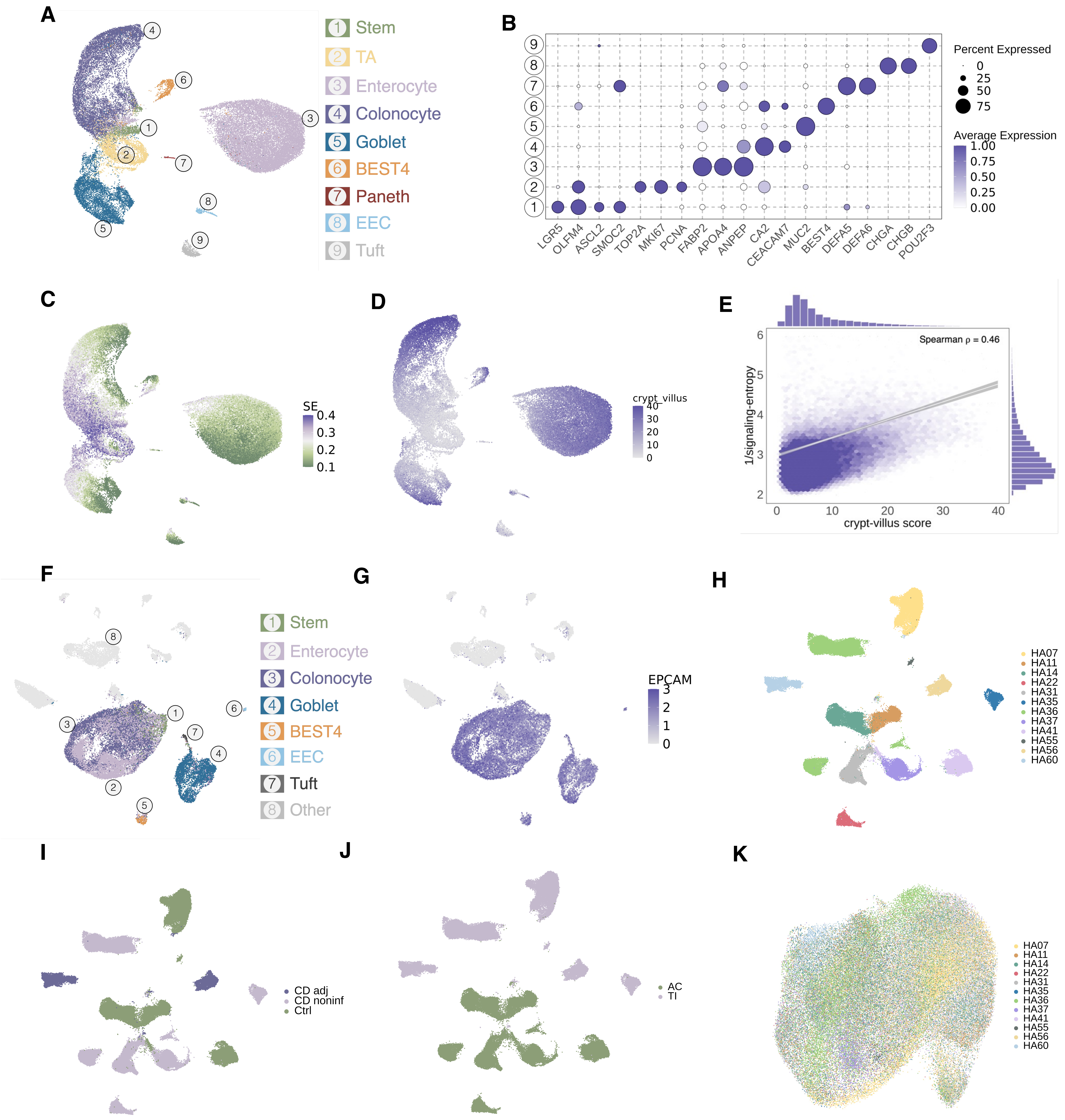

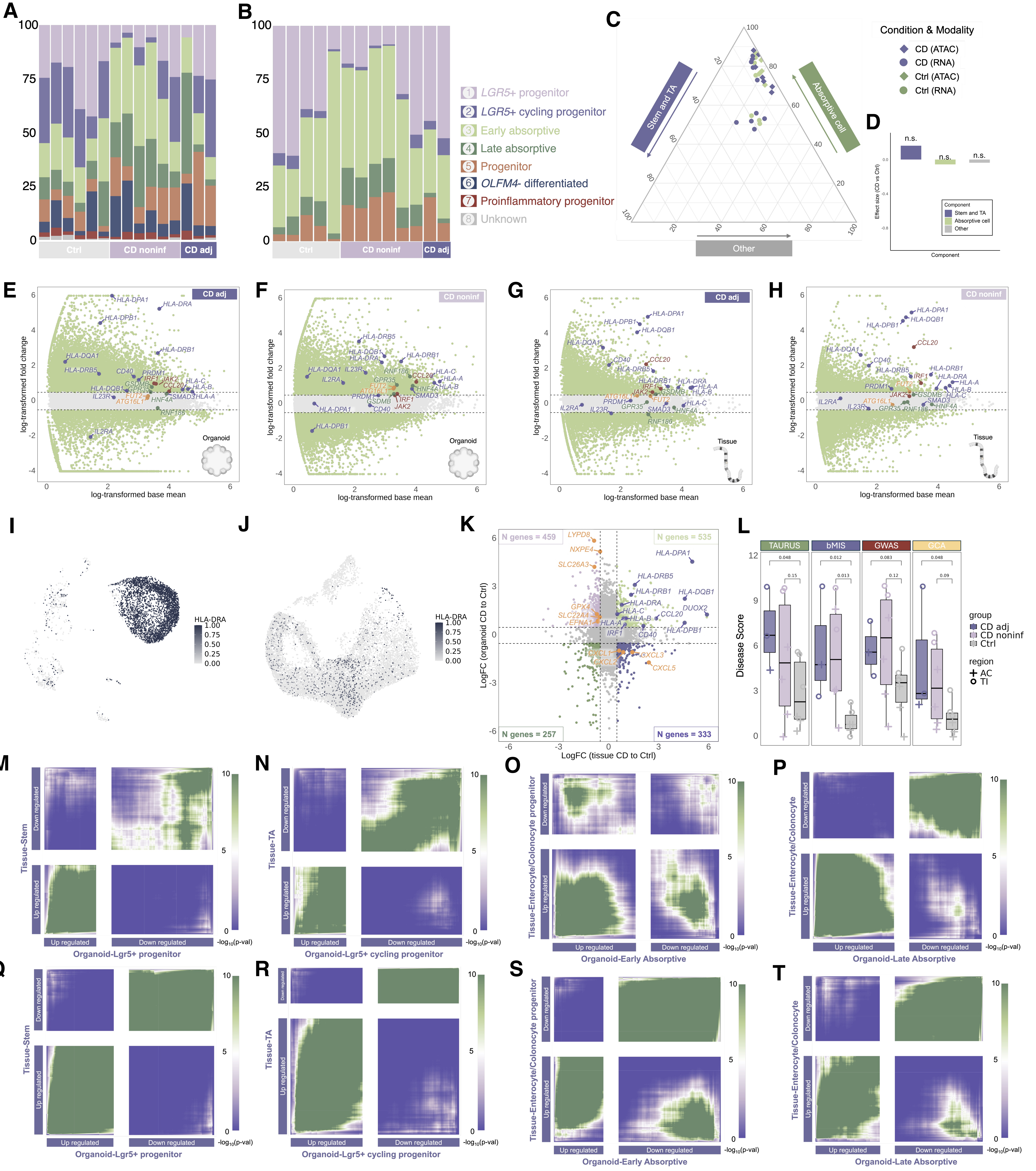

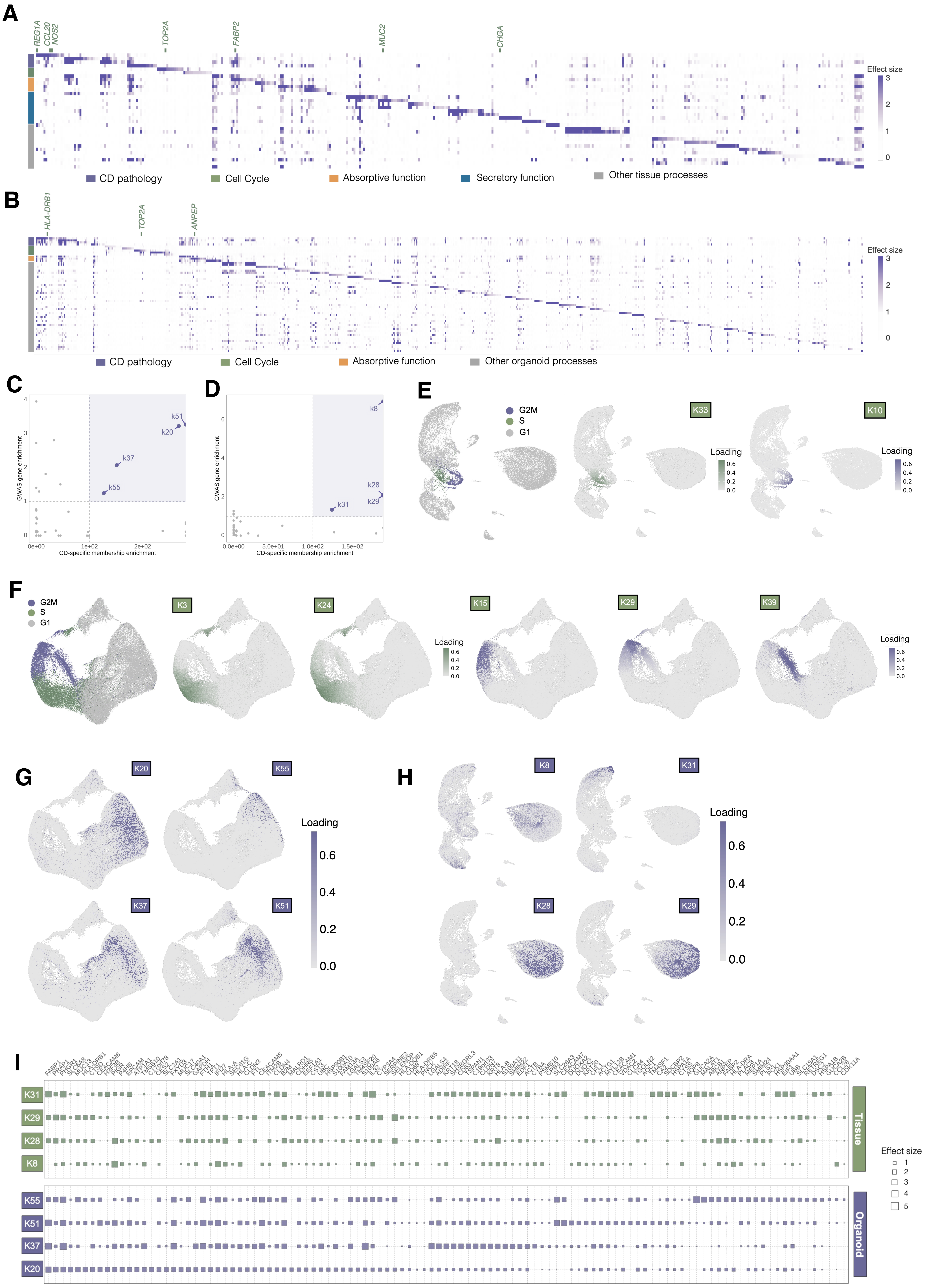

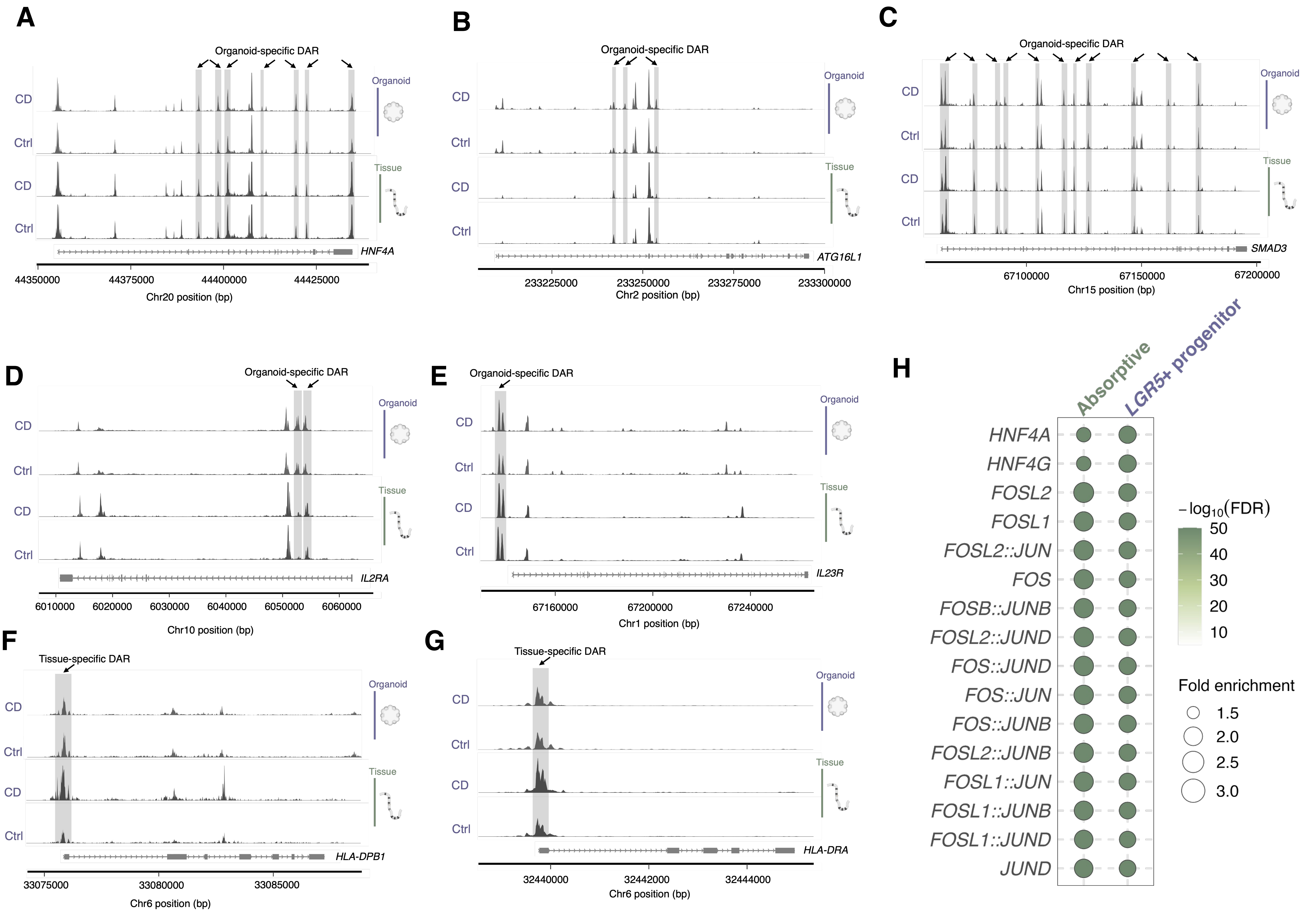

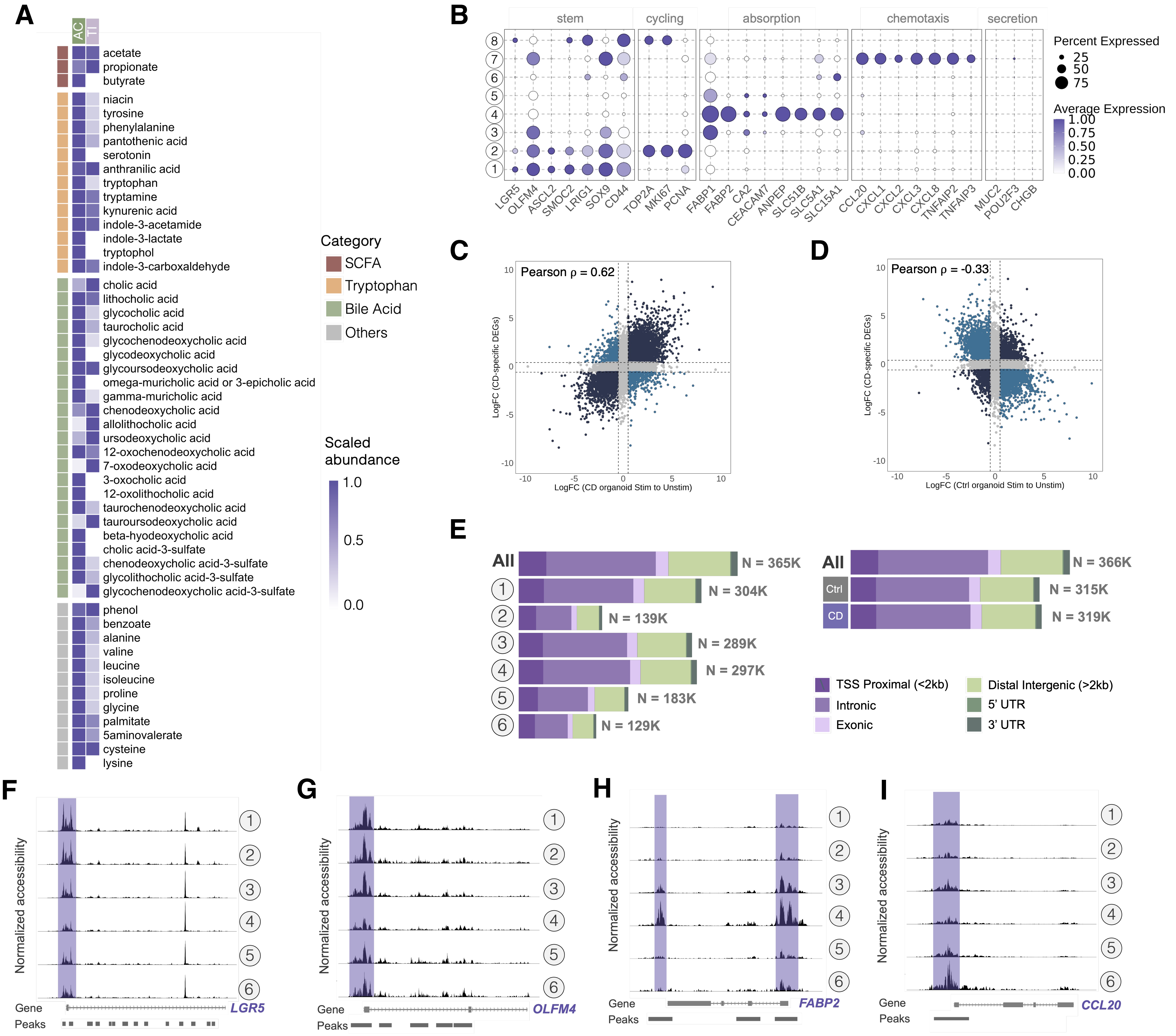
